# Unification of Sparse Bayesian Learning Algorithms for Electromagnetic Brain Imaging with the Majorization Minimization Framework

**DOI:** 10.1101/2020.08.10.243774

**Authors:** Ali Hashemi, Chang Cai, Gitta Kutyniok, Klaus-Robert Müller, Srikantan S. Nagarajan, Stefan Haufe

**Affiliations:** Uncertainty, Inverse Modeling and Machine Learning Group, Technische Universität Berlin, Germany; Machine Learning Group, Technische Universität Berlin, Germany; Berlin Center for Advanced Neuroimaging (BCAN), Charité – Universitätsmedizin Berlin, Germany; Institut für Mathematik, Technische Universität Berlin, Germany; Department of Radiology and Biomedical Imaging, University of California, San Francisco, CA, USA; National Engineering Research Center for E-Learning, Central China Normal University, China; Mathematisches Institut, Ludwig-Maximilians-Universität München, Germany; Department of Physics and Technology, University of Tromsø, Norway; BIFOLD – Berlin Institute for the Foundations of Learning and Data, Berlin, Germany; Department of Brain and Cognitive Engineering, Korea University, Seoul, South Korea; Max Planck Institute for Informatics, Saarbrücken, Germany; Mathematical Modelling and Data Analysis Department, Physikalisch-Technische Bundesanstalt Braunschweig und Berlin, Germany; Bernstein Center for Computational Neuroscience, Berlin, Germany

**Keywords:** Electro-/Magnetoencephalography, Brain Source Imaging, Type I/II Bayesian Learning, Non-convex, Majorization-Minimization, Noise Learning, Hyperparameter Learning

## Abstract

Methods for electro- or magnetoencephalography (EEG/MEG) based brain source imaging (BSI) using sparse Bayesian learning (SBL) have been demonstrated to achieve excellent performance in situations with low numbers of distinct active sources, such as event-related designs. This paper extends the theory and practice of SBL in three important ways. First, we reformulate three existing SBL algorithms under the *majorization-minimization* (MM) framework. This unification perspective not only provides a useful theoretical framework for comparing different algorithms in terms of their convergence behavior, but also provides a principled recipe for constructing novel algorithms with specific properties by designing appropriate bounds of the Bayesian marginal likelihood function. Second, building on the MM principle, we propose a novel method called *LowSNR-BSI* that achieves favorable source reconstruction performance in low signal-to-noise-ratio (SNR) settings. Third, precise knowledge of the noise level is a crucial requirement for accurate source reconstruction. Here we present a novel principled technique to accurately learn the noise variance from the data either jointly within the source reconstruction procedure or using one of two proposed cross-validation strategies. Empirically, we could show that the monotonous convergence behavior predicted from MM theory is confirmed in numerical experiments. Using simulations, we further demonstrate the advantage of LowSNR-BSI over conventional SBL in low-SNR regimes, and the advantage of learned noise levels over estimates derived from baseline data. To demonstrate the usefulness of our novel approach, we show neurophysiologically plausible source reconstructions on averaged auditory evoked potential data.

## I. Introduction

Electro- and Magnetoencephalography (EEG/MEG) are non-invasive techniques for measuring brain electrical activity with high temporal resolution. As such, both have become indispensable tools in basic neuroscience and clinical neurology. The downside of both techniques, however, is that their sensors are located far away from the neural generators of the measured brain electrical activity. EEG/MEG measurements are therefore characterized by low spatial resolution and highly overlapping contributions of multiple brain sources in each sensor. The mathematical model of the EEG/MEG sensing procedure can be described by the linear *forward model*

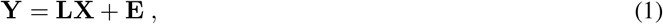

which maps the electrical activity of the brain sources, **X**, to the sensor measurements, **Y**. The *measurement matrix* **Y** ∈ ℝ^*M*×*T*^ captures the activity of *M* sensors attached at different parts of the scalp at *T* time instants, **y**(*t*) ∈ ℝ^*M*×1^, *t* = 1, …, *T*, while the *source matrix*, **X** ∈ ℝ^*N*×*T*^, consists of the unknown activity of *N* brain sources located in the cortical gray matter at the same time instants, **x**(*t*) ∈ ℝ^*N*×1^, *t* = 1, …, *T*. The matrix **E** = [**e**(1), …, **e**(*T*)] ∈ ℝ^*M*×*T*^ represents *T* time instances of independent and identically distributed (i.i.d.) zero mean white Gaussian noise with variance *σ*^2^, 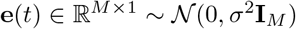, *t* = 1, …, *T*, which is assumed to be independent of the source activations. The linear forward mapping from **X** to **Y** is given by the *lead field* matrix **L** ∈ ℝ^*M*×*N*^, which is here assumed to be known. In practice, **L** can be computed using discretization methods such as the Finite Element Method (FEM) for a given head geometry and known electrical conductivities using the quasi-static approximation of Maxwell’s equations [1]–[4].

The goal of brain source imaging (BSI) is to infer the underlying brain activity **X** from the EEG/MEG measurement **Y** given the lead field matrix **L**. Unfortunately, this inverse problem is highly ill-posed as the number of sensors is typically much smaller than the number of locations of potential brain sources. Thus, a unique solution cannot be found without introducing further mathematical constraints or penalties, which are often referred to as regularizers. In addition, the leadfield matrix is typically highly ill-conditioned even for small numbers of sensors, introducing numerical instabilities in the inverse estimates.

Interestingly, regularization can also be interpreted in a Bayesian framework, where the regularizer introduces prior knowledge or assumptions about the nature of the true sources into the estimation [5], [6]. A common assumption is that the number of active brain sources during the execution of a specific mental task is small, i.e., that the spatial distribution of the brain activity is sparse. This assumption can be encoded in various ways. Classical approaches [7] employ super-Gaussian prior distributions to identify solutions in which most of brain regions are inactive. In these approaches Maximum-a-Posteriori (MAP) estimation, also termed *Type-I learning*, is used. Later work [8] has shown that hierarchical Bayesian models achieve better reconstructions of sparse brain signals by employing a separate Gaussian prior for each brain location. The variances at each location are treated as unknown (hyper-) parameters, which are estimated jointly with the source activity. This approach is called Sparse Bayesian Learning (SBL), Type-II Maximum-Likelihood (Type-II ML) estimation or simply *Type-II learning* [9]–[11].

Type-II learning generally leads to non-convex objective functions, which are non-trivial to optimize. A number of iterative algorithms have been proposed [8]–[10], [12]–[14], which, due to employing distinct parameter update rules, differ in their convergence guarantees, rates and overall computational complexity. Being derived using vastly different mathematical concepts such as fixed point theory and expectation-maximization (EM), it has, however, so far been difficult to explain the observed commonalities and differences, advantages and disadvantages of Type-II methods in absence of a common theoretical framework, even if the properties of individual algorithms have been extensively studied [12].

The primary contribution of this paper is to introduce *Majorization-Minimization (MM)* ([15], [16], and references therein) as a flexible algorithmic framework within which different SBL approaches can be theoretically analyzed. Briefly, MM is a family of iterative algorithms to optimize general non-linear objective functions. In a minimization setting, MM replaces the original cost function in each iteration by an upper bound, or majorization function, whose minimum is usually easy to find. The objective value at the minimum is then used to construct the bound for the following iteration, and the procedure is repeated until a local minimum of the objective is reached. Notably, MM algorithms are popular in many disciplines in which Type-II learning problems arise, such as, e.g., telecommunications [17]–[21] and finance [22], [23]. The concept of MM is, however, rarely explicitly referenced in EEG/MEG brain source imaging, even though it has been used implicitly [24], [25]. We demonstrate here that three popular SBL variants, denoted as *EM*, *MacKay*, and *convex-bounding based* SBL, can be cast as majorization-minimization methods employing different types of upper bounds on the marginal likelihood. This view as variants of MM helps explain, among other things, the guaranteed convergence of these algorithms to a local minimum. The characteristics of the chosen bounds determine the reconstruction performance and convergence rates of the resulting algorithms. The MM framework additionally offers a principled way of constructing new SBL algorithms for specific purposes by designing appropriate bounds.

Therefore, a second contribution of this paper is the development of a new SBL algorithm, called LowSNR-BSI, that is especially suitable for low signal-to-noise ratio (SNR) regimes. Real-world applications of EEG/MEG brain source imaging are often characterized by low SNR, where the power of unwanted noise sources can be comparable to the power of the signal of interest. This holds in particular for the reconstruction of ongoing as well as induced (non-phase-locked) oscillatory activity, where no averaging can be performed prior to source reconstruction. Current SBL algorithms may suffer from reduced performance in such low-SNR regimes [26]–[28]. To overcome this limitation, we propose a novel MM algorithm for EEG/MEG source imaging, which employs a bound on the SBL cost function that is particularly suitable for low-SNR regimes.

As a third contribution, this paper discusses principled ways to estimate the sensor noise variance *σ*^2^, which is assumed to be known in the first part of the paper. Determining the goodness-of-fit of the optimal model, the value of this variable exerts a strong impact on the overall reconstruction [29]. Technically being another model hyperparameter, the noise variance is, however, rarely estimated as part of the model fitting. Instead, it is often determined prior to the model fitting from a baseline recording. This approach can, however, lead to suboptimal results in practice or be even inapplicable, e.g., when resting state data are analyzed. Here we present a number of alternatives to estimate the noise variance in Type-I and Type-II brain source imaging approaches. Building on work by [30], we derive an analytic update rule, which enables the adaptive estimation of the noise variance within various SBL schemes. Moreover, we propose two novel cross-validation (CV) schemes from the machine learning field to determine the noise variance parameter.

We conduct extensive ground-truth simulations in which we compare LowSNR-BSI with popular source reconstruction schemes including existing SBL variants, and in which we systematically study the impact of different strategies to estimate the noise level *σ*^2^ from the data.

The outline of the paper is as follows: In Section II, a comprehensive review of Type-II BSI methods is presented. In Section III, we unify the Type-II methods described in Section II within the MM framework, and in Section IV, we derive LowSNR-BSI algorithm within the same framework. Section V introduces numerous principled ways for estimating the sensor noise variance. Simulation studies, real data analysis, and discussions are presented in Sections VI, VII, and VIII, respectively. Finally, Section IX concludes the paper.

## II. Bayesian Learning

The ill-posed nature of the EEG/MEG inverse problem can be overcome by assuming a prior distribution *p*(**X**) for the source activity. The posterior distribution of the sources after observing the data **Y**, *p*(**X**|**Y**), is given by Bayes’ rule:

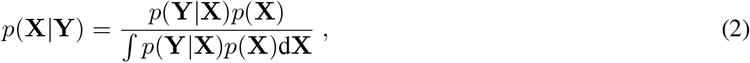

where the conditional probability *p*(**Y**|**X**) in the numerator denotes the *likelihood*, while the term in the denominator, *∫ p*(**Y**|**X**)*p*(**X**)d**X** = *p*(**Y**), is referred to as *model evidence* or *marginal likelihood*. However, note that the posterior is often not analytically tractable, as evaluating the integral in the model evidence is intractable for many choices of prior distributions and likelihoods.

### Remark 1.

*Priors fulfill the same practical purpose as regularizers even if they are motivated from a different perspective, e.g., the Bayesian formalism, in this paper. Besides, we regard the Bayesian perspective as a helpful technical vehicle to inspire and generate flexible priors for shaping more plausible solutions.*

### A. Type-I Bayesian Learning

As the model evidence in Eq. (2) only acts as a scalar normalization for the posterior, its evaluation can be avoided if one is only interested in the most probable source configuration **X** rather than the full posterior distribution. This point estimate is known as the maximum-a-posteriori (MAP) estimate:

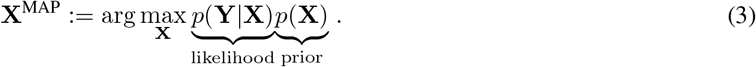

Assuming i.i.d. Gaussian sensor noise, the likelihood reads:

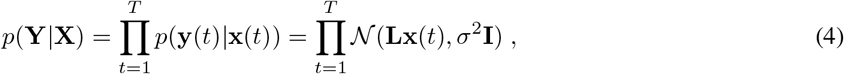

and the resulting MAP estimate (3) is given by

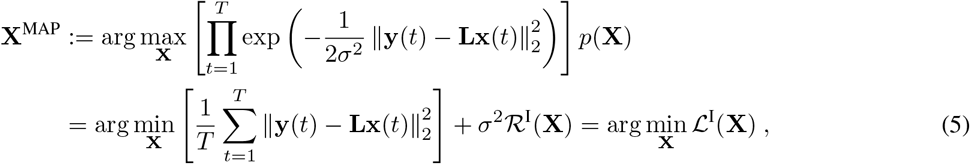

where 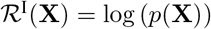 and 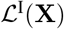 denotes the Bayesian Type-I learning (MAP) objective function.

Note that this expression can be interpreted as a trade-off between two optimization goals, where the first (log-likelihood) term in (5) penalizes model errors using a quadratic loss function and the second (log-prior) term penalizes deviations of the solution from the assumed spatial or temporal properties of the brain sources encoded in 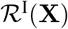. The trade-off between these two optimization goals is defined by the ratio of the noise variance *σ*^2^ and the variance of the prior distribution. As the latter is hardly known in practice, a *regularization parameter λ* ∝ *σ*^2^ subsuming both variables is introduced, which can be tuned to adjust the relative importance of both penalties in the optimization.

Several existing algorithms are characterized by different choices of a prior. For instance, choosing a Gaussian prior distribution leads to the classical minimum-norm estimate [31]–[33], which also goes by the names 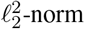 (or Tikhonov) regularization and “ridge regression” in the statistics and machine learning literature. The choice of a Laplace prior leads to the minimum-current estimate [7], which is also known as *ℓ*_1_-norm regularization or “LASSO” regression. Besides, hierarchical Bayesian priors with automatic depth weighting have been used to infer brain activity from EEG/MEG data [34]. More complex priors have been also used to incorporate anatomical information of the sources [35]–[37] or to encode assumptions on the spatial, temporal and/or spectral structure of the sources. Respective methods include FOCUSS [38], S-FLEX [39], [40], MxNE [41], ir-MxNE [42], TF-MxNE [43], irTF-MxNE [44], and STOUT [45], which all enforce sparsity in different domains such as Gabor frames or cortical patches through appropriate norm constraints.

### B. Type-II Bayesian Learning

While in the MAP approach the prior distribution is fixed, it is sometimes desirable to consider entire families of distributions *p*(**X**|***γ***) parameterized by a set of hyper-parameters *γ*. These hyper-parameters can be learned from the data along with the model parameters using a hierarchical empirical Bayesian approach [9]–[11]. In this maximum-likelihood Type-II (ML-II, or simply Type-II) approach, *γ* is estimated through the maximum-likelihood principle:

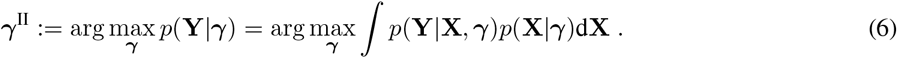

Computation of the conditional density *p*(**Y**|***γ***) is formally achieved by integrating over all possible source distributions **X** for any given choice of ***γ***. The maximizer of Eq. (6) then determines a data-driven prior distribution *p*(**X**|***γ***^II^). Plugged into the MAP estimation framework Eq. (3), this gives rises to the Type-II source estimate **X**^II^.

As the conditional density *p*(**Y**|***γ***) for a given ***γ*** is identical to the model evidence in Eq. (2), this approach also goes by the name evidence maximization [14], [30]. Concrete instantiations of this approach have further been introduced under the names *sparse Bayesian learning* (SBL) [10] or *automatic relevance determination* (ARD) [46], *kernel Fisher discriminant* (KFD) [9], *variational Bayes* (VB) [12], [47] and iteratively-reweighted MAP estimation [13], [38]. Interested readers are referred to [48] for a comprehensive survey on Bayesian machine learning techniques for EEG/MEG signals. To distinguish all these Type-II variants from classical ML and MAP approaches not involving hyperparameter learning, the latter are also referred to as Type-I approaches.

#### Remark 2.

*The marginal likelihood formulation in Type-II Bayesian learning,* *Eq.* (6), *enables estimation of flexible priors with many parameters from data. This stands in contrast to the use of classical cross-validation techniques to learn hyperparameters of regularizers, which works for very few parameters only (in most cases only a single scalar regularization constant).*

### C. Sparse Bayesian Learning and Champagne

A Type-II estimation framework with particular relevance for EEG/MEG source imaging is SBL. In this framework, the *N* modeled brain sources are assumed to follow independent univariate Gaussian distributions with zero mean and distinct unknown variances 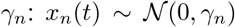, *n* = 1, …, *N*. In the SBL solution, the majority of variances is zero, thus effectively inducing spatial sparsity of the corresponding source activities. Such sparse solutions are physiologically plausible in task-based analyses, where only a fraction of the brain’s macroscopic structures is expected to be consistently engaged. This consideration has led [11] to propose the *Champagne* algorithm for brain source imaging, which is rooted in the concept of SBL. Compared to Type-I approaches achieving sparsity through *ℓ*_1_-norm minimization, Champagne has shown significant performance improvement with respect to EEG/MEG source localization [8], [27].

Just as most existing approaches, Champagne makes the simplifying assumption of statistical independence between time samples. This leads to the following expression for the distribution of the sources:

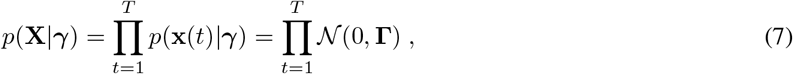

where ***γ*** = [*γ*_1_, …, *γ_N_*]^⊤^ and **Γ** = diag(***γ***). Note that, in task-based analyses, the noise variance *σ*^2^ can be estimated from a baseline (resting state) recording. In the first part of this paper, it is, therefore, assumed to be known.

#### Remark 3.

*Electrophysiological data are known to possess a complex intrinsic autocorrelation structure. Here, we consider priors that make the simplifying assumption of independence between time samples, which is consistent with most existing works in the field [7], [31], [32], [39], [41]. Importantly, using such simplifying priors generally does not prevent the resulting inverse solutions to have time structure. Nevertheless, priors modeling the known properties of the latent variables more accurately might lead to better reconstructions especially in low-sample regimes. Preliminary work shows that priors modeling temporal structure with autoregressive models can indeed improve the reconstruction of autocorrelated source [24].*

The parameters of the SBL model are the unknown sources as well as their variances. As computation of the integral in Eq. (6) is infeasible, Champagne considers an approximation, where the variances *γ_n_*, *n* = 1, …, *N*, are optimized based on the current estimates of the sources in an alternating iterative process. Given an initial estimate of the variances, the posterior distribution of the sources is a Gaussian of the form [8], [49, Chapter 4]

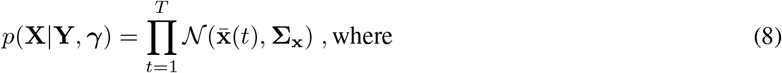

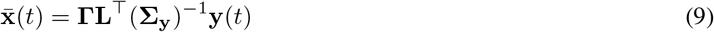

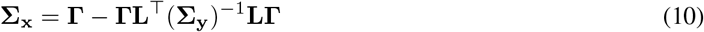

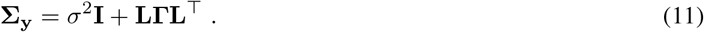

The estimated posterior parameters 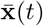 and **Σ_x_** are then in turn used to update the estimate of the variances *γ_n_, n* = 1, …, *N* as the minimizer of the negative log of the marginal likelihood *p*(**Y**|***γ***), which is given by [8]:

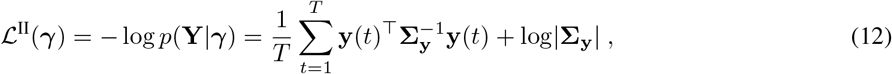

where | · | denotes the determinant of a matrix. This process is repeated until convergence. Given the final solution of the hyperparameter ***γ***^II^, the point estimate **x**^II^ of the source activity is obtained from the posterior mean of the estimated source distribution: 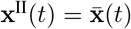. Note that given the definition of the empirical sample covariance matrix as 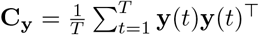, the term 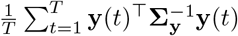 in Eq. (12) can be rewritten as 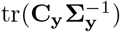, so that Eq. (12) becomes [8, Section II]

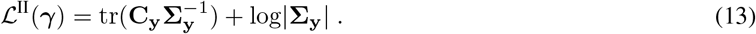

Note that, in this form, the loss function Eq. (13) bears an interesting similarity to the *log-determinant (log-det) Bregman divergence* in information geometry [50]. This perspective on Type-II loss function enables a common viewpoint for Type-I and Type-II methods.

By invoking mathematical tools based on Legendre-Fenchel duality theory, the cost function Eq. (12) can be formulated equivalently as another cost function, 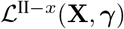, whose optimizers, {***γ***\*, **X**\*}, are derived by performing a joint minimization over **X** and ***γ*** [14, see also Section II-B], [51], [52]:

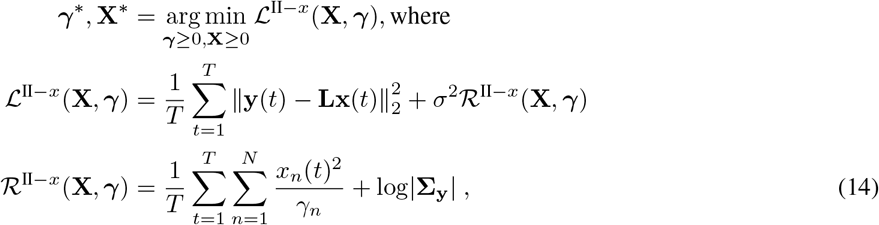

where 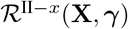 denotes a regularizer that depends on the data, **x**(*t*), and where *x_n_*(*t*) denotes the activity of source *n* at time instant *t*. Then, as each source *x_n_*(*t*) is also a function of *γ_n_* according to Eq. (9), the term 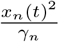 goes to zero when *γ_n_* → 0.

#### Remark 4.

*In contrast to standard MAP estimation, the effective priors obtained within our hierarchical Bayesian framework, e.g.,* 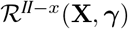 *in* *Eq.* (14), *are not fixed. They depend on parameters that can be tuned and learned from the data; thus, Type-II priors have the ability and flexibility to capture the actual properties of the observed real data.*

We will use the formulation in Eq. (14) to derive alternative optimization schemes for Champagne in Sections II-C2 and II-C3.

#### 1) EM Champagne

As the cost function Eq. (12) is non-convex in ***γ***, the quality of the obtained solution depends substantially on the properties of the employed numerical optimization algorithm. Crucially, algorithms might not only differ with respect to their convergence properties but may also lead to different solutions representing distinct local minima of Eq. (12). The first algorithm for mimimizing Eq. (12) has been introduced by [12] and is an application of the expectation-maximization (EM) formalism [53]. As can be shown, Eqs. (9)–(11) correspond to the expectation (E) step of the EM algorithms with respect to the posterior distribution *p*(**X**|**Y**, ***γ***). The maximization (M) step of the EM formalism with respect to ***γ*** then leads to the update rule

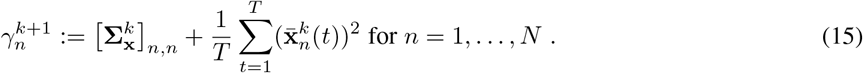

Final estimates of both parameters are obtained by iterating the updates (9)–(11) and (15) until convergence. The resulting algorithm is known as the EM variant of the Champagne algorithm [49, Chapter 4] [12] in the field of brain source imaging.

#### 2) Convex-bounding based Champagne

As the EM algorithm outlined above has been shown to have slow convergence speed, alternative minimization strategies have been proposed. Two such variants, a convex-approximation based approach and the so-called MacKay update, have been proposed in [12] and further practically investigated in [27]. Considering that the log-determinant in Eq. (14) is concave, the convex-bounding based variant of Champagne constructs a linear upper bound based on the concave conjugate of log |*σ*^2^**I** + **LΓL**^⊤^|, defined as *w*\*(**z**),

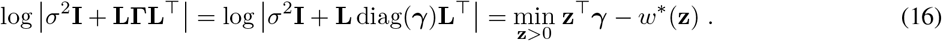

With this upper bound, and for a fixed value of ***γ***, the auxiliary variable **z** can be derived as the tangent hyperplane of the log|**Σ_y_**|:

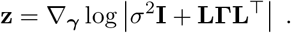

Note that the concave conjugate is obtained as a result of applying Legendre-Fenchel duality theory (see, e.g., [51], [52]) on the concave function log |*σ*^2^**I** + **LΓL**^⊤^| as follows: 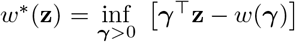, where *w*(***γ***) = log |*σ*^2^**I** + **LΓL**^⊤^| denotes our target concave function.

By inserting Eq. (16) instead of log|**Σ_y_**| into Eq. (14), the non-convex penalty function Eq. (14) is replaced by the convex function

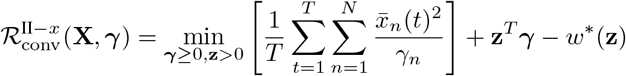

in each step of the optimization. The final estimates of **X**, ***γ*** and **z** are obtained by iterating between following update rules until convergence:

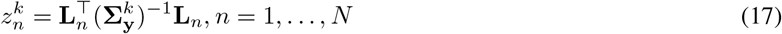

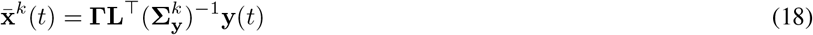

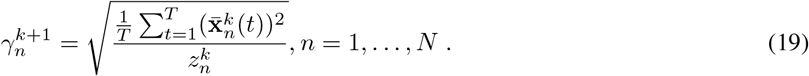

Here, **L**_*n*_ in (17) denotes the *n*-th column of the lead field matrix.

#### 3) MacKay Update for Champagne

The MacKay update proposed in [12, Section III.A-2] can be derived in a similar fashion as the convex-bounding based update using different auxiliary functions and variables. By defining new variables *κ_n_* ≔ log(*γ_n_*) for *n* = 1, …, *N*, the non-convex term log |*σ*^2^**I**+ **L**diag(***γ***)**L**^⊤^| in Eq. (16) can be written as:

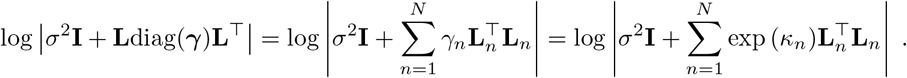

Then, one can introduce another surrogate function [12, Appendix-B]

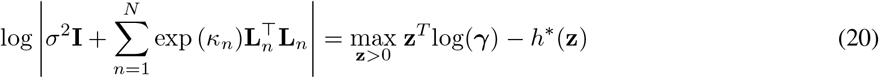

for the log |*σ*^2^**I** + **LΓL**^⊤^|, where *h*\*(**z**) denotes the convex conjugate of 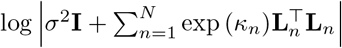 in contrast to the concave conjugate counterpart, *w*\*(**z**) used in Eq. (16). Substituting (20) into Eq. (14) leads to a so-called *min-max optimization program* for optimizing the non-convex penalty function 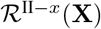, which alternates between minimizations over ***γ*** and maximizations of the bound in (20):

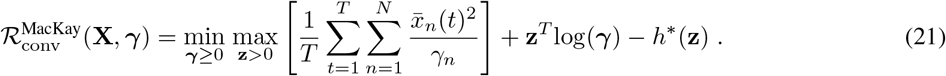

Let 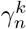 denote the value of *γ_n_* in the *k*-th iteration. Inserting ***γ***^*k*^ into Eq. (21) and minimizing with respect to 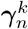 requires that the derivatives

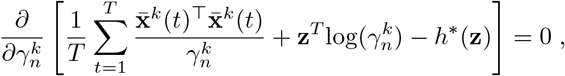

for *n* = 1, …, *N*, vanish. The resulting function is then maximized with respect to **z** [12, Appendix-B], which leads to the so-called MacKay update for optimizing Eq. (14) [12, Section A-2]:

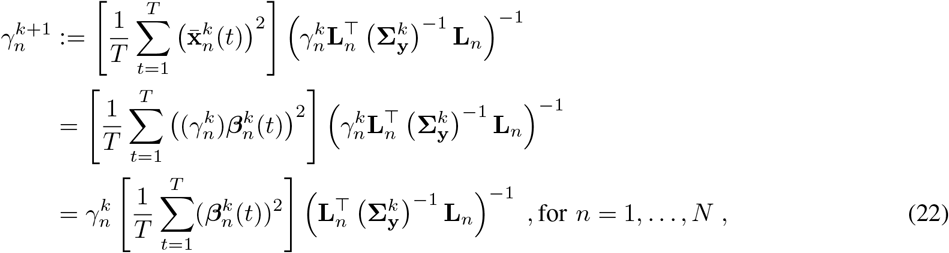

where 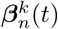 is defined as follows: 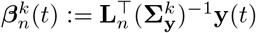 for *n* = 1, …, *N*.

## III. Unification of Sparse Bayesian Learning algorithms with the Majorization-Minimization (MM) Framework

In this section, we first briefly review theoretical concepts behind the MM algorithmic framework [15], [54]–[56]. Then, we formally characterize Champagne variants as MM algorithms by suggesting upper bounds on the cost function Eq. (14) that, when employed within the MM framework, yield the same update rules as the original algorithms. The first three rows of Table I list the update rules and mathematical formalism used in this section.

**TABLE I.** This table summarizes the update rules presented in Section II-C and their corresponding MM upper-bounds that will be utilized in Sections III and IV.

|  | Update Rule | Mathematical Formalism Used | Inequality Used |
| --- | --- | --- | --- |
| EM | $\gamma_n^{k+1} := [\mathbf{\Sigma}_{\mathbf{x}}^k]_{n,n} + \frac{1}{T} \sum_{t=1}^T (\bar{\mathbf{x}}_n^k(t))^2$ | Expectation-Maximization Formalism | Jensen's Inequality |
| Convex Bounding | $\gamma_n^{k+1} := \sqrt{\left[\frac{1}{T} \sum_{t=1}^T (\bar{\mathbf{x}}_n^k(t))^2\right]} \left(\mathbf{L}_n^\top (\mathbf{\Sigma}_{\mathbf{y}}^k)^{-1} \mathbf{L}_n\right)^{-1/2}$ | Concave Conjugate | Taylor Expansion |
| MacKay | $\gamma_n^{k+1} := \left[\frac{1}{T} \sum_{t=1}^T (\bar{\mathbf{x}}_n^k(t))^2\right] \left(\gamma_n^k \mathbf{L}_n^\top (\mathbf{\Sigma}_{\mathbf{y}}^k)^{-1} \mathbf{L}_n\right)^{-1}$ | Change of Variable + Convex Conjugate | Taylor Expansion |
| LowSNR-BSI | $\gamma_n^{k+1} := \sqrt{\left[\frac{1}{T} \sum_{t=1}^T (\bar{\mathbf{x}}_n^k(t))^2\right]} \left(\mathbf{L}_n^\top \mathbf{L}_n\right)^{-1/2}$ | MM Principle in Low-SNR Setting | Taylor Expansion |

### A. Majorization-Minimization

Majorization-minimization is a promising strategy for solving general non-linear optimization programs. Compared to other popular optimization paradigms such as (quasi)-Newton methods, MM algorithms enjoy guaranteed convergence to a stationary point [16]. The MM class covers a broad range of common optimization algorithms such as *proximal methods* and *convex-concave procedures (CCCP)* [16, Section IV], [57], [58]. While such algorithms have been applied in various contexts, such as non-negative matrix factorization [59] and massive MIMO systems for wireless communication [19], [21], their advantages have so far rarely been made explicit in the context of brain source imaging [24], [25], [60].

We define an original optimization problem with the objective of minimizing a continuous function *f* (**u**) within a closed convex set 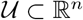:

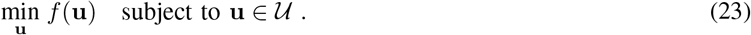

Then, the idea of MM can be summarized as follows. First, construct a continuous *surrogate function g*(**u**|**u**^*k*^) that upper-bounds, or *majorizes*, the original function *f* (**u**) and coincides with *f* (**u**) at a given point **u**^*k*^:

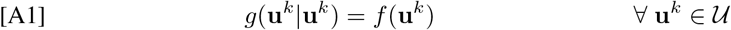

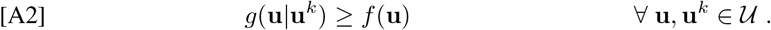

Second, starting from an initial value **u**^0^, generate a sequence of feasible points **u**^1^, **u**^2^, …, **u**^*k*^, **u**^*k*+1^ as solutions of a series of successive simple optimization problems, where

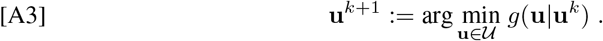

Note that the performance of MM algorithms heavily depends on the choice of a suitable surrogate function, which should, on one hand, faithfully reflect the behavior of the original non-convex function Eq. (23) while, on the other hand, be easy to minimize.

#### Definition 1.

*Any algorithm fulfilling conditions [A1]–[A3] is called a Majorization Minimization (MM) algorithm.*

#### Corollary 1.

*An MM algorithm has a* descending trend *property, whereby the value of the cost function f decreases in each iteration: f* (**u**^*k*+1^) ≤ *f* (**u**^*k*^).

*Proof.* The proof is included in Appendix B.

While Corollary 1 guarantees a descending trend, convergence requires additional assumptions on particular properties of *f* and *g* [54], [55]. For the smooth functions considered in this paper, we require that the derivatives of the original and surrogate functions coincide at **u**^*k*^:

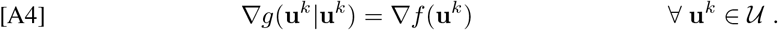

Then, the following, stronger, theorem holds.

#### Theorem 1.

*For an MM algorithm that additionally satisfies [A4], every limit point of the sequence of minimizers generated through [A3] is a stationary point of the original optimization problem* *Eq.* (23).

*Proof.* A detailed proof can be found in [54, Theorem 1].

Note that since we are working with smooth functions, conditions [A1]–[A4] are sufficient to prove convergence to a stationary point according to Theorem 1 (see [15], [54], [56] and [53], [61]) for proofs of the convergence behaviour of other MM algorithms such as expectation maximization.

#### Remark 5.

*Corollary 1 implies that if a surrogate function is constructed to fulfill conditions [A1] and [A2], and if the next feasible point of the algorithm is always assigned as the minimizer of the surrogate function based on [A3], the resulting* MM *algorithm decreases f* (**u**) *in each step. Although a weaker condition than [A3], i.e., g*(**u**^*k*+1^|**u**^*k*^) ≤ *g*(**u**^*k*^|**u**^*k*^), *is sufficient for a descending trend, we only consider MM algorithms in this paper; thus, condition [A3] is a crucial requirement. As we have shown in Theorem 1, [A3] is further required to prove guaranteed convergence of an MM algorithm.*

We now show that three algorithms that have been proposed for solving the SBL cost function Eq. (12) can all be cast as instances of the MM framework invoking different majorization functions on 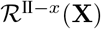. For the convex-bounding based approach as well as the algorithm using MacKay updates, the full set of conditions [A1]–[A4] in Theorem 1 are proven. Due to the considerations made above, we, however, only prove Corollary 1 for the EM-based Champagne algorithm.

#### 1) EM Update as MM

It is known that the EM algorithm is a special case of MM framework using Jensen’s inequality to construct the surrogate function [16], [56]. Here, we work out the specific surrogate function for the SBL cost function Eq. (12) (i.e., the negative log marginal likelihood).

As Wipf and Nagarajan have shown [12, Section III.A-1], the EM algorithm for Type-II problems consists of the following two parts: For the E-step, the posterior *p*(**X**|**Y**, ***γ**^k^*) is obtained given the value of ***γ*** at *k*-th iteration, ***γ***^*k*^. The M-step then solves:

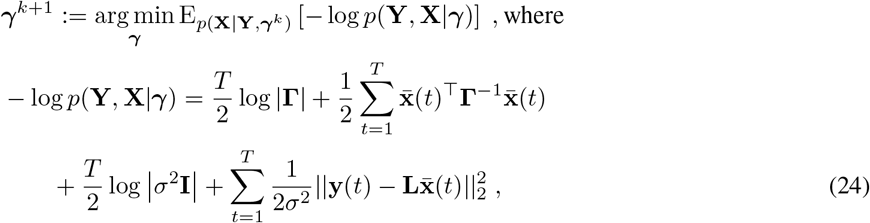

which leads to the update rule in Eq. (15).

##### Proposition 1.

*The EM based Champagne algorithm is an MM algorithm fulfilling Corollary 1, where the negative log-likelihood loss*, −log *p*(**Y**|***γ***), *is majorized by the following surrogate function*

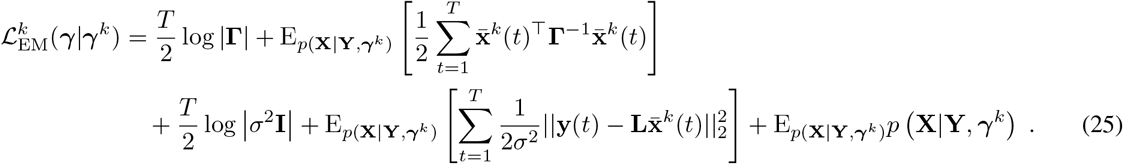

*Proof.* A detailed proof can be found in Appendix C.

Note that the *EM* algorithm is also equivalent to the restricted maximum likelihood (ReML) [62] and dynamic statistical parametric mapping (dSPM) approaches [63] for solving the sparse EEG/MEG inverse problem, which, thereby, can also be interpreted as instances of minimization-majorization.

#### 2) Convex-bounding Based Approach as MM

We start by recalling the non-convex penalty 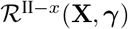 as defined in Eq. (14):

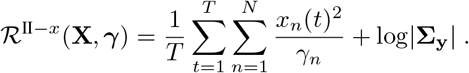

By setting 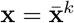 to the value obtained by the convex-bounding based method in the *k*-th iteration, the following holds:

##### Proposition 2.

*The convex-bounding based Champagne algorithm is an MM algorithm fulfilling Theorem 1, where* 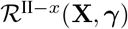 *is majorized by the following surrogate function:*

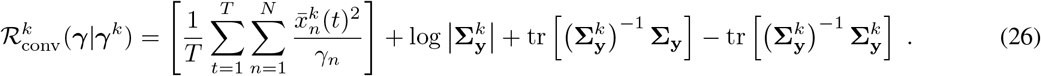

*Proof.* A detailed proof is provided in Appendix D.

#### 3) MacKay Update as MM

Similar to convex-bounding, we can show that the Mackay updates for Champagne can be viewed as an MM algorithm.

##### Proposition 3.

*The Champagne variant employing MacKay updates is an MM algorithm fulfilling Theorem 1, where* 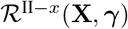 *is majorized by* 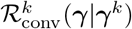.

*Proof.* The proof is similar to that of Proposition 2 and provided in Appendix E.

To summarize this section, we have shown that three popular strategies for solving the SBL problem in Eq. (12), namely the EM, the MacKay, and the convex bounding based approaches, can be characterized as MM algorithms. Importantly, this perspective provides a common framework for comparing different Champagne algorithms. For example, we can derive and compare certain characteristics of Champagne algorithms directly based on the properties of the majorization functions they employ. Conversely, it is also possible to design specific majorization functions that are optimal in a specific sense, leading to new source reconstruction algorithms.

## IV. LowSNR-Brain Source Imaging (LowSNR-BSI)

Here, we assume a low-SNR regime, as it is common in BSI applications. SNR is defined in sensor space as signal power, 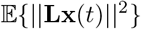, divided by noise power, 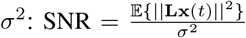, and can be expressed in dB scale as SNR_dB_ = 10log10 (SNR). In many practical applications, we are interested in solving the BSI problem for SNR_dB_ ≤ 0; that is, when the noise power is comparable to the power of the signal or even larger. Although the algorithms presented in Sections III-A1–III-A3 achieve satisfactory performance in terms of computational complexity, their reconstruction performance degrades significantly in low-SNR regimes. This behavior has been theoretically shown in [26, Section VI-E] and has also been confirmed in several simulation studies [27], [28].

In order to improve the performance of SBL in low-SNR settings, we propose a novel MM algorithm by constructing a surrogate function for Eq. (12) specifically for this setting. Based on [19], we propose the following convex surrogate function:

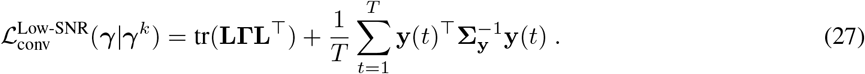

The following proposition is based on results in [19].

### Proposition 4.

*The surrogate function* *Eq.* (27) *majorizes the Type-II loss function* *Eq.* (12) *and results in an MM algorithm that fulfills Theorem 1. For SNR* → 0, *Eq.* (12) *converges to* *Eq.* (27):

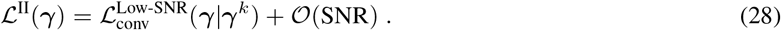

*Proof.* A detailed proof of this result is presented in Appendix F. The main idea is to first normalize the sensor and source covariance matrices by *σ*^2^ and then consider the eigenvalue decomposition of tr(**LΓL**^⊤^) as **LΓL**^⊤^ = **UPU**^⊤^ with **P** = diag(*p*_1_, …, *p_M_*). These two steps result in the following equality: log |**Σ_y_**| = log |**I**+ **UPU**^⊤^|. Finally, the proof is completed by leveraging the concavity of the log(·) function and using a Taylor expansion around the eigenvalues of **LΓL**^⊤^, i.e., *p_i_*, for *i* = 1, …, *M*.

Note that, as a result of Proposition 4, the behaviour of the non-convex SBL cost function Eq. (12) is more and more well approximated in the vicinity of the current estimate by the proposed surrogate function Eq. (27) as the noise level increases, which sets it apart from existing surrogate functions. Therefore, the proposed bound is particularly suitable in low-SNR regimes.

In contrast to the original SBL cost function Eq. (12), the surrogate function Eq. (27) is convex and has unique minimum that can be found analytically in each iteration of the optimization. To find the optimal value of ***γ*** = [*γ*_1_,…, *γ_N_*]^⊤^, we first take the derivative of (27) with respect to each *γ_n_* for *n* = 1,…, *N*, and then set it to zero, which yields the following closed-form solution for ***γ*** = [*γ*_1_,…, *γ_N_*]^⊤^:

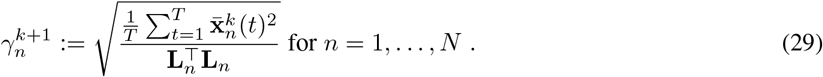

A detailed derivation of Eq. (29) can be found in Appendix G. We call the algorithm obtained by iterating between (9)–(11) and (29) *LowSNR-Brain Source Imaging (LowSNR-BSI)*. In practice, values exactly equal to zero may not be obtained for the *γ_n_*. Therefore, an *active-set* strategy is employed. Given a threshold *γ*_thresh_, those variances *γ_n_* for which *γ_n_* < *γ*_thresh_ holds are set to zero in each iteration of the algorithm. Algorithm 1 summarizes the steps of LowSNR-BSI. Table I allows for a direct comparison of the LowSNR-BSI update rule (last column) and the corresponding update rules of other Champagne variants derived within the MM framework.

## V. Automatic Estimation of the Noise Level

### A. Adaptive Noise Learning

It is common practice to estimate the noise variance *σ*^2^ from baseline data prior to solving the EEG/MEG inverse problem [64]–[70]. However, a baseline estimate may not always be available or may not be accurate enough, say, due to inherent non-stationarities in the data/experimental setup. Here, we argue that estimating the noise parameter from the to-be-reconstructed data can significantly improve the reconstruction performance even compared to a baseline estimate. To this end, we here derive data-driven update rules that allow us to tune estimate the noise variance, *σ*^2^ within the source reconstruction procedure using the Champagne and LowSNR-BSI algorithms, where we build on prior work by [9], [10], [30], [71], [72]. Practically we introduce the shortcut *λ* = *σ*^2^ to underscore that *λ* is a tunable parameter whose estimate can substantially deviate from the baseline estimate in practice. We then treat *λ* as another model hyperparameter, similar to the source variances *γ_n_*. Thus, in each step of learning cycles of the Champagne and LowSNR-BSI algorithms, we also minimize the loss function 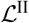 with respect to *λ*, where the remaining parameters **Γ** and **Σ_x_** are fixed to the values obtained in the preceding iteration. This leads to the following theorem:

**Algorithm 1:**
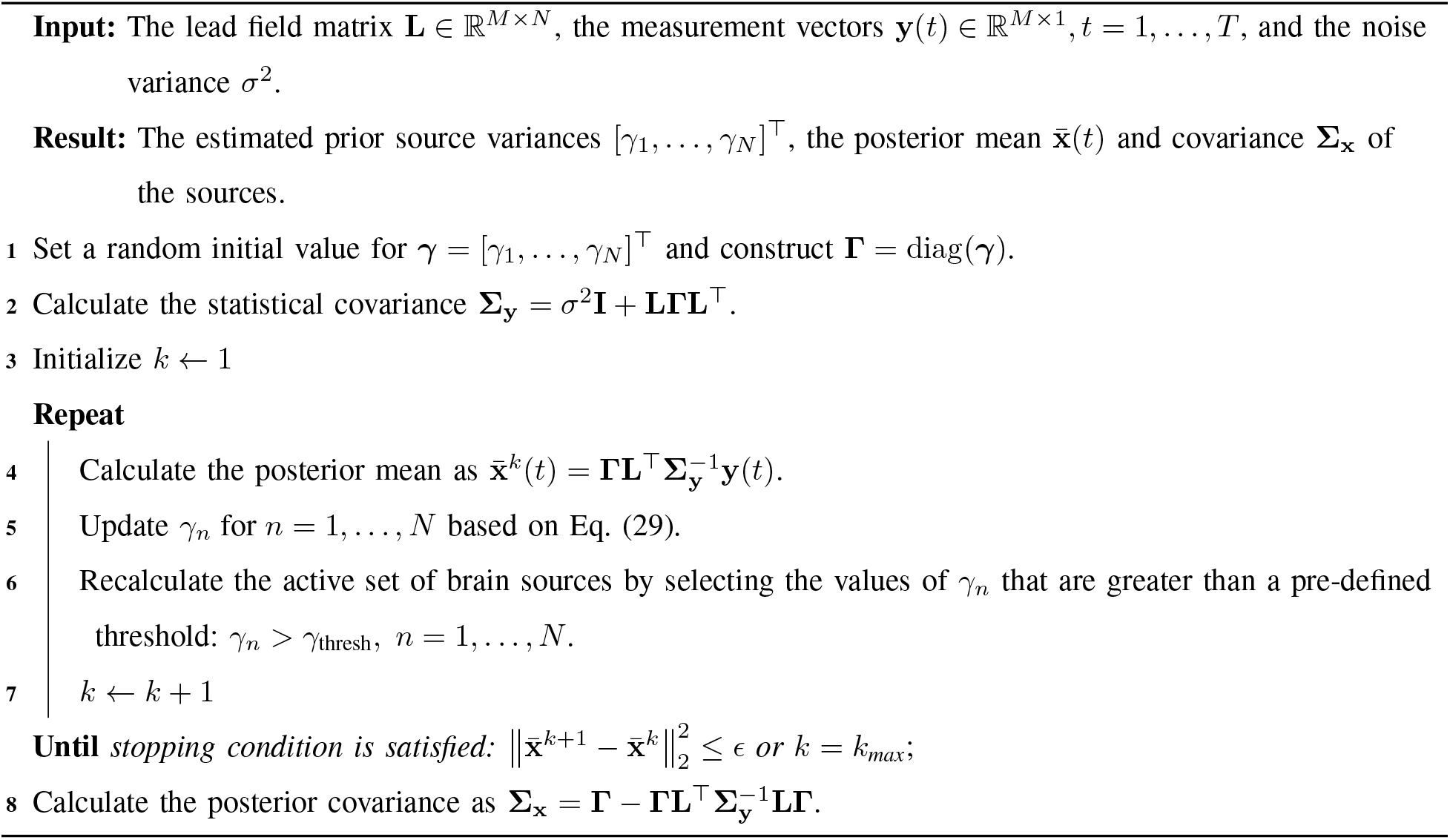
LowSNR-BSI algorithm.

#### Theorem 2.

*The minimization of* 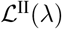 *with respect to λ,*

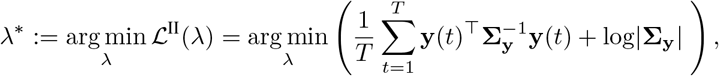

*yields the following update rule for λ at the* (*k* + 1)*-th iteration, assuming* **Γ**^*k*^ *and* 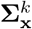 *be fixed values obtained in the* (*k*)*-th iteration*:

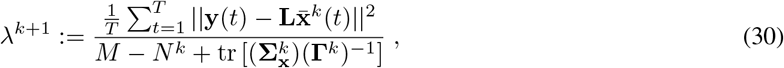

*where N^k^ denotes the number of non-zero voxels identified at iteration k through an active-set strategy*.

*Proof.* A detailed proof can be found in Appendix H.

As shown in Algorithm (1), our implementation uses an active-set strategy that only selects the non-zero voxels at each iteration based on a threshold. Therefore, at the initial steps of the algorithm, *N^k^* = *N* since all source variances are initialized randomly. But, when the algorithm proceeds, the number of non-zero voxels decreases as a result of our active-set strategy, which results in smaller values for *N^k^*.

### B. Cross-validation Strategies

In the previous section, we proposed to estimate the noise variance *λ* = *σ*^2^ *in-sample* such that the SBL likelihood according to Eq. (12) was maximized, which led to an analytic update rule. As, under our assumption of homoscedastic sensor noise, *λ* is only a single scalar parameter, it moreover becomes feasible make use of robust model selection techniques employing the concept of cross-validation (CV), whose aim it is to maximize the *out-of-sample* likelihood [73]–[75]. To this end, the data are split into two parts. On the so-called *training set*, the model parameters is fitted for a wide range of possible values of *λ*, which are fixed within each individual optimization. The likelihoods of the fitted models are then evaluated on the held-out data parts, called the *test sets*. The choice of *λ* that maximizes the empirical likelihood on the test data is then used as an unbiased estimate of the noise variance. It is well-known from the field of machine learning that cross-validation effectively overcomes the problem of model overfitting in small samples. Here, we introduce two CV strategies employing different ways of splitting the data.

#### 1) Temporal Cross-validation

In temporal CV, the temporal sequence of the data samples is split into *k* different contiguous blocks (folds) [76], [77]. Here, we use *k* = 4. Three folds form the training set, 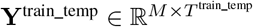, on which we fit the Champagne and LowSNR-BSI models for a range of *λ*s. On the remaining fold, 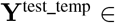 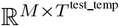, the Type-II log-likelihood (c.f. Eqs. (12) and (13))

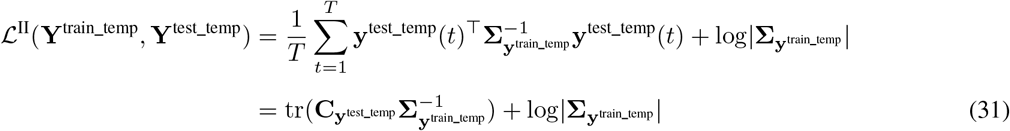

is then evaluated. Note that in Eq. (31) the model covariance 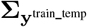 that has been determined on the training data **Y**^train_temp^ is combined with the empirical covariance of the hold-out data **Y**^test_temp^, which were not used during model fitting. Thus, Eq. (31) is the *out-of-sample* Type-II log-likelihood. It has been theoretically shown [26], [78] that the Type-II log-likelihood function is a metric on the second-order information of the sensors closely related to the log-det Bregman divergence (discrepancy) between statistical (model) and empirical covariances [50], [79]. The choice of *λ* that minimizes that discrepancy on hold-out data is, therefore, a sensible estimate for the true noise variance. We provide further details on the relation between the SBL likelihood and the log-det Bregman divergence in Appendix A.

#### 2) Spatial Cross-validation

In spatial CV, the data are not split into temporal segments but by dividing the available EEG/MEG sensors into the training and test sets. This variant has been proposed by [29], [40]. Here, we again use *k* = 4 folds, where we randomly assign 75% of the sensors to the training set, 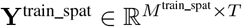, and the remaining 25% to the test set, 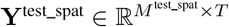. On the training sensors, Champagne and LowSNR-BSI are fitted using the corresponding portion of the leadfield matrix, **L**^train_spat^, for the same range of *λ*s as used in temporal CV. The sources, **X**^train_spat^ ∈ ℝ^*N*×*T*^, estimated from the fitted models are then mapped back to the sensor space, and the out-of-sample Type-I log-likelihood (c.f. Eq. (5)) is evaluated on the hold-out (test) sensors:

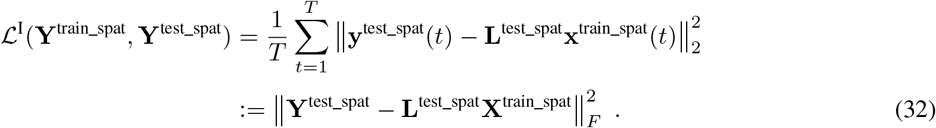

Note that, while the Type-II log-likelihood has an interpretation as a Bregman divergence between model and empirical covariance matrices, the Type-I log-likelihood is the Frobenius norm or mean-squared error (MSE) 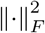 of the model residuals, i.e., the average squared Euclidean distance between empirical and modeled observation vectors. Thus, while the Type-II likelihood compares model and observations in terms of their second-order statistics, the Type-I likelihood uses only first-order information. As in temporal CV, the value of *λ* that minimizes the MSE on the test sensors is selected as the final noise estimate.

## VI. Simulations

We conducted an extensive set of simulations, in which we compared the reconstruction performance of the proposed LowSNR-BSI algorithm to that of Champagne and two additional widely-used source reconstruction schemes for a range of different SNRs. We also tested impact of the proposed noise learning schemes (adaptive, temporal CV and spatial CV) on the source reconstruction performance compared to estimating the noise level from baseline data.

### A. Pseudo-EEG Signal Generation

#### Forward Modeling

Populations of pyramidal neurons in the cortical gray matter are known to be the main drivers of the EEG signal [80]. Here, we use a realistic volume conductor model of the human head to model the linear relationship between primary electrical source currents in these populations and the scalp surface potentials captured by EEG electrodes. The New York Head model [4] provides a segmentation of an average human head into six different tissue types. In this model, 2004 dipolar current sources were placed evenly on the cortical surface and 58 sensors were placed on the scalp according to the extended 10-20 system [81]. In accordance with the predominant orientation of pyramidal neuron assemblies, the orientation of all source currents was fixed to be perpendicular to the cortical surface, so that only scalar source amplitudes needed to be estimated. Finite-element modeling was used to compute the lead field matrix, **L** ∈ ℝ^58×2004^, which serves as the forward model in our simulations.

#### Source Generation

We simulated a sparse set of *N*_0_ = 3 active sources, which were placed at random positions on the cortex. The temporal activity of each source was generated by a univariate linear autoregressive (AR) process, which models the activity at time *t* as a linear combination of the *P* past values:

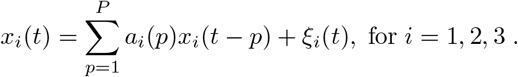

Here, *a_i_*(*p*) for *i* = 1, 2, and 3 are linear AR coefficients, and *P* is the order of the AR model. The model residuals *ξ_i_*(·) for *i* = 1, 2 and 3 are also referred to as the innovation process; their variance determines the stability of the overall AR process. We here assume uncorrelated standard normal distributed innovations, which are independent for all sources. In the following, we use stable AR systems of order *P* = 5.

#### Noise Model

To simulate the electrical neural activity of the underlying brain sources, *T* = 20 data points were sampled from the AR process described above. Corresponding dipolar current sources were then placed at random locations, yielding sparse source activation vectors **x**(*t*). Source activations **X** = [**x**(1), …, **x**(*T*)] were mapped to the 58 EEG sensors through application of the lead field matrix **L**:

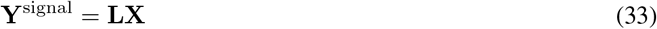

Next, we added Gaussian white noise to the sensor-space signal. To this end, noise was randomly sampled from a standard normal distribution and normalized with respect to its Frobenius norm. A weighted sum of signal and noise contributions then yielded the pseudo-EEG signal

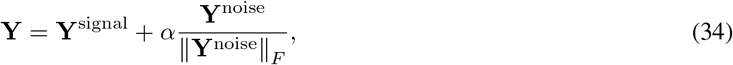

where *α* determines the signal-to-noise ratio in sensor space. For a given *α*, the noise variance is obtained as *σ*^2^ = 1/*M* tr[**Σ_e_**], for 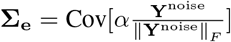, and the SNR (in dB) is calculated as 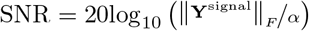. Since our goal is to investigate the effect of noise variance estimation on the performance of the proposed algorithms, we fixed the noise variance in each set of simulations so as to obtain distributions of performance metrics for a number of similar effective SNR values. We conducted four sets of simulations using *α* = {2, 1.5, 1, 0.5}, corresponding to average noise variances of *σ*^2^ = {37.4 × 10^−3^, 21.0 × 10^−3^, 9.4 × 10^−3^, 2.3 × 10^−3^} and average SNRs of SNR = {0.33, 2.17, 4.87, 11.40} (dB). Each set of simulations consists of 100 experiments, in which source locations and time series as well as noise realizations were randomly sampled.

In addition to the pseudo-EEG signal, a pseudo baseline measurement containing only noise but no signal was generated. The sole purpose of this measurement was to provide an empirical estimate of the noise variance as a baseline for our joint source reconstruction and noise estimation approaches, which estimate the same quantity from the summed pseudo-EEG signal. To ensure sufficiently precise baseline estimation, 300 noise samples were generated, normalized, and scaled by *α* as in Eq. (34) for each experiment.

### B. Source reconstruction

We applied Champagne and LowSNR-BSI to the synthetic datasets described above. The variances of all voxels were initialized randomly by sampling from a standard normal distribution. The optimization programs were terminated either after reaching convergence (defined by a relative change of the Frobenius-norm of the reconstructed sources between subsequent iterations of less than 10^−8^), or after reaching a maximum of *k*_max_ = 3000 iterations.

In each experiment, we evaluated the algorithms using 40 predefined choices of the noise variance ranging from *λ* = ⅓*σ*^2^ to *λ* = 30*σ*^2^. In addition, *λ* was estimated from data using the techniques introduced in Section V. We observed that the variance estimated from baseline data, 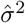 (averaged over all EEG channels) was typically almost identical to the ground-truth value *λ* = *σ*^2^ used to simulate the data. The reconstruction performance obtained using this value was therefore included in the comparison as a baseline. Performance at baseline noise level was compared to the performance obtained using adaptive learning of the noise using Eq. (30) as well as using spatial or temporal cross-validation. Note that, for temporal CV, we generated *T* = 80 samples, so that we obtained 60 samples in each training set and 20 samples in each test fold. Due to the increased number of training samples, this method, therefore, has an advantage over the remaining ones. For spatial CV, due to the spatial blur introduced by volume conduction, there is a limit on how focal the measured sensor-space electrical potentials or magnetic fields can be, and the signal will usually be distributed over all sensors. Therefore, a setting in which all ‘signal-carrying’ electrodes will end up either in the training or test set is unlikely to occur in practice. Using, for example, *k* = 4 random splits, it is ensured that the training set will typically capture the signal pattern well. The test set in this approach is only used to evaluate the out-of-sample likelihood on the remaining sensors, while no model fitting needs to take place. Therefore, missing certain aspects of the signal pattern in the test set does not pose a critical problem, especially if multiple splits are conducted.

#### Remark 6.

*The fact that real M/EEG data have time structure is acknowledged in our simulation setting by modeling source time courses as AR processes. The resulting samples of the training and test sets thereby become dependent. Technically, this violates the i.i.d. assumption underlying the theory of CV. However, one can argue that training and test sets are de-facto independent since the leakage from one set to another is small compared to the length of the data. In the spatial CV approach, in contrast, the sensors of the training and test sets are strongly dependent on another, because of the spatial blur introduced by volume conduction. Nevertheless, as we observe in Sections VI-D and VII, spatial CV works very well both in simulations and real data analysis. This observation suggests that the cross-validation approach can work even if the i.i.d. assumption is violated, in line with previous literature [29], [40], [74], [82].*

In addition to Champagne and LowSNR-BSI, two non-SBL source reconstruction schemes were included for comparison. As an example of a sparse Type-I method based on *ℓ*_1_-norm minimization, S-FLEX [40] was used. As spatial basis functions, unit impulses were used, so that the resulting estimate was identical to the so-called minimum-current estimate [7]. In addition, the eLORETA estimate [33], a smooth inverse solution based on weighted 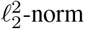 minimization was used. eLORETA was used with 5% regularization, whereas S-FLEX was fitted so that the residual variance was consistent with the ground-truth noise level. Note that the 5% rule is chosen as it gives the best performance across a subset of regularization values ranging between 0.5% to 15%.

### C. Evaluation Metrics

Source reconstruction performance was evaluated according to the following metrics. First, the *earth mover’s distance* (EMD, [39], [83]) was used to quantify the spatial localization accuracy. The EMD metric measures the cost needed to transform two probability distributions, defined on the same metric domain, into each other. It was applied here to the *N* × 1 amplitude distributions of the true and estimated sources, which were obtained by taking the voxel-wise *ℓ*_2_-norm along the time domain. EMD scores were normalized to be in [0, 1]. Second, the error in the reconstruction of the source time courses was measured. To this end, Pearson correlation between all pairs of simulated and reconstructed (i.e., those with non-zero activations) sources was measured. Each simulated source was matched to a reconstructed source based on maximum absolute correlation. Time course reconstruction error was then defined as one minus the average of these absolute correlations across sources. Finally, the runtime of the algorithms was measured in seconds (*s*).

### D. Results

Figure 1 shows the EMD (upper row), the time course reconstruction error (middle row) and the negative log-likelihood loss value (lower low) incurred by Champagne and LowSNR-BSI for two SNR settings (SNR = 0.33 dB and SNR = 11.40 dB). Four different schemes of estimating the noise level from data (estimation from baseline data, adaptive learning, spatial CV, and temporal CV) are compared. Note that we found previously that the ground-truth noise variance *λ* = *σ*^2^ used in the simulation is generally accurately estimated from baseline data, which is referred to as ‘baseline’ in the figure, 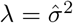. Interestingly, however, this baseline is optimal only for LowSNR-BSI, and only with respect to temporal source reconstruction. For Champagne, and with respect to the spatial source reconstruction performance of LowSNR-BSI, the choice of the baseline noise variance turns out to be suboptimal, as it is outperformed by all three proposed schemes that estimate the noise variance from the actual (task) data to be reconstructed (‘Adaptive Learning’, ‘Spatial CV’ and ‘Temporal CV’). Interestingly, noise levels estimated using Spatial CV lead to near-optimal reconstruction performance in a broad variety of settings, in line with observations made in [29], [40]. All proposed noise learning schemes converge to points in the vicinity of the minimum of the SBL loss function Eq (12).

**Fig. 1.**
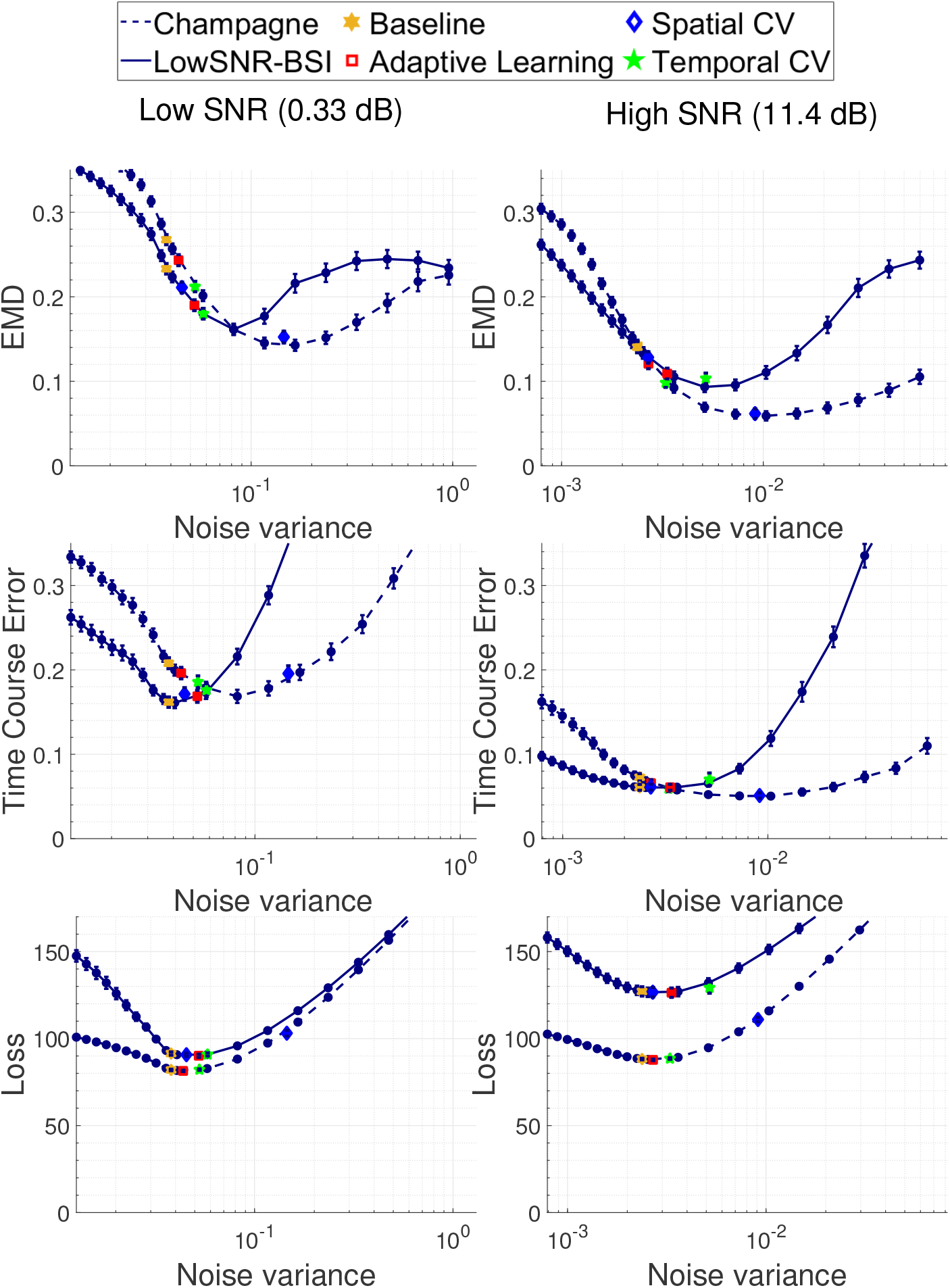
Source reconstruction performance of Champagne and LowSNR-BSI in two different SNR regimes (low SNR: 0.33 dB, left column; high SNR: 11.4 dB, right column). Spatial reconstruction error is measured in terms of the earth-mover’s distance, and is shown in the upper row, while time course reconstruction error is shown in the middle row. The lower row demonstrates the negative log-likelihood loss, SBL loss function Eq. (12), incurred by Champagne and LowSNR-BSI algorithms.

The EMD in our setting only depends on the spatial distribution of the sources. Therefore, the EMD is not able to fully capture potential advantages resulting from modeling temporal characteristics of the correlated EEG/MEG time courses. As a result, it is not highly aligned with the values of the loss. This explains the observed discrepancies between the loss function and the EMD values of Champagne and LowSNR-BSI in Figure 1. To assess the reconstruction of the temporal characteristics of the brain sources, we also measure the time course error. All four variants of LowSNR-BSI algorithms not only outperform their Champagne counterparts but also approach the minimal achievable time course error. High EMD performance of Champagne with Spatial CV does not lead to high performance in terms of time course error as well as regarding the negative log-likelihood loss. For all algorithms, regularization values resulting in a smaller EMD metric can be found. However, this observation does not imply a practical benefit of any algorithm as the ground-truth is unknown in real-world situations.

Figure 2 further compares the source reconstruction performance of the four noise estimation variants separately for Champagne and LowSNR-BSI for a range of four SNR values. As already observed in Figure 1, all three proposed approaches for noise variance estimation (adaptive learning, spatial CV, and temporal CV) lead to better source reconstruction performance than the estimation from baseline data. Overall, spatial CV for Champagne and temporal CV for LowSNR-BSI achieve the best combination of spatial and temporal reconstruction performance.

**Fig. 2.**
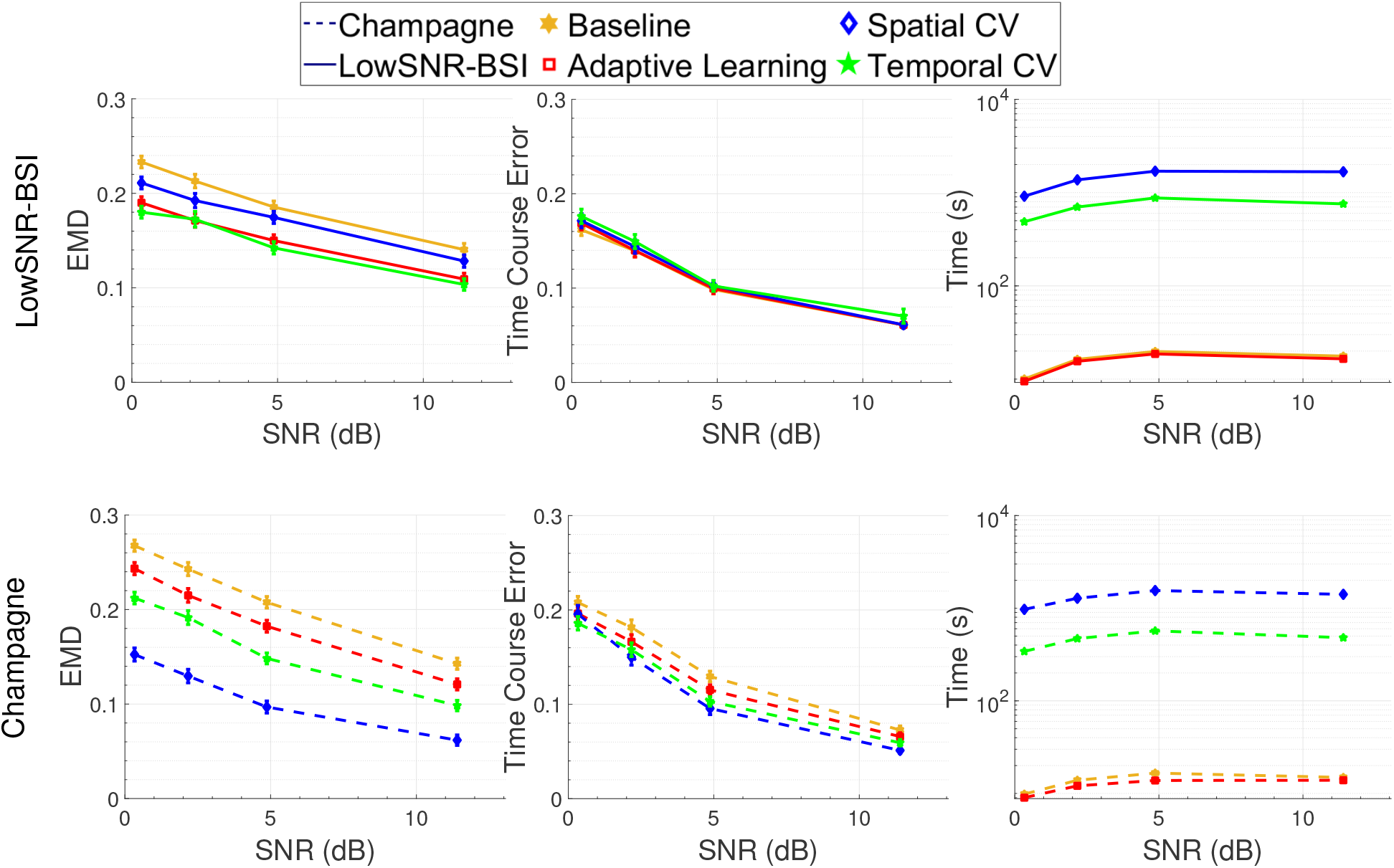
Source reconstruction performance of four different variants of LowSNR-BSI (upper row) and Champagne (lower row). The noise variance was estimated from baseline data (ground truth), using adaptive learning, or using spatial or temporal cross-validation. Performance was evaluated for four SNRs (SNR = {0.33, 2.17, 4.87, 11.40} dB) and with respect to three different metrics (spatial reconstruction according to the earth-mover’s distance – left column, time course reconstruction error – middle column, and computational complexity according to the runtime (in seconds) – right column).

The superior performance of CV techniques, however, comes at the expense of higher computational complexity of the source reconstruction. As Figure 2 demonstrates, using CV techniques with the specified numbers of folds increases the runtime of Champagne and LowSNR-BSI by approximately two orders of magnitude (10^3^*s* ~ 10^4^*s*) compared to the runtimes of eLORETA, S-FLEX, and the baseline and adaptive learning variants of Champagne and LowSNR-BSI (1*s* ~ 10*s*).

Figure 3 provides an alternative depiction of the data presented in Figure 2, which allows for a more direct comparison of Champagne and LowSNR-BSI. As benchmark algorithms, eLORETA [33] and S-FLEX [40] are also included in the comparison. It can be seen that LowSNR-BSI in the baseline mode, using adaptive noise learning, and using temporal CV consistently outperforms Champagne in terms of spatial localization accuracy, in particular in low-SNR settings. This behavior indeed confirms the advantage of the surrogate function, 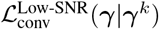, which is designed to provide a better approximation of the non-convex SBL cost function in low-SNR regimes, as presented in Section IV. Consequently, as the SNR decreases, the gap between LowSNR-BSI and Champagne further increases. In terms of the time course reconstruction error, LowSNR-BSI shows a similar improvement over Champagne when the SNR is low. However, the magnitude of this improvement is not as pronounced as observed for the EMD metric. The only setting in which Champagne consistently outperforms LowSNR-BSI is when spatial CV is used to estimate the noise variance, and spatial reconstruction performance is evaluated.

**Fig. 3.**
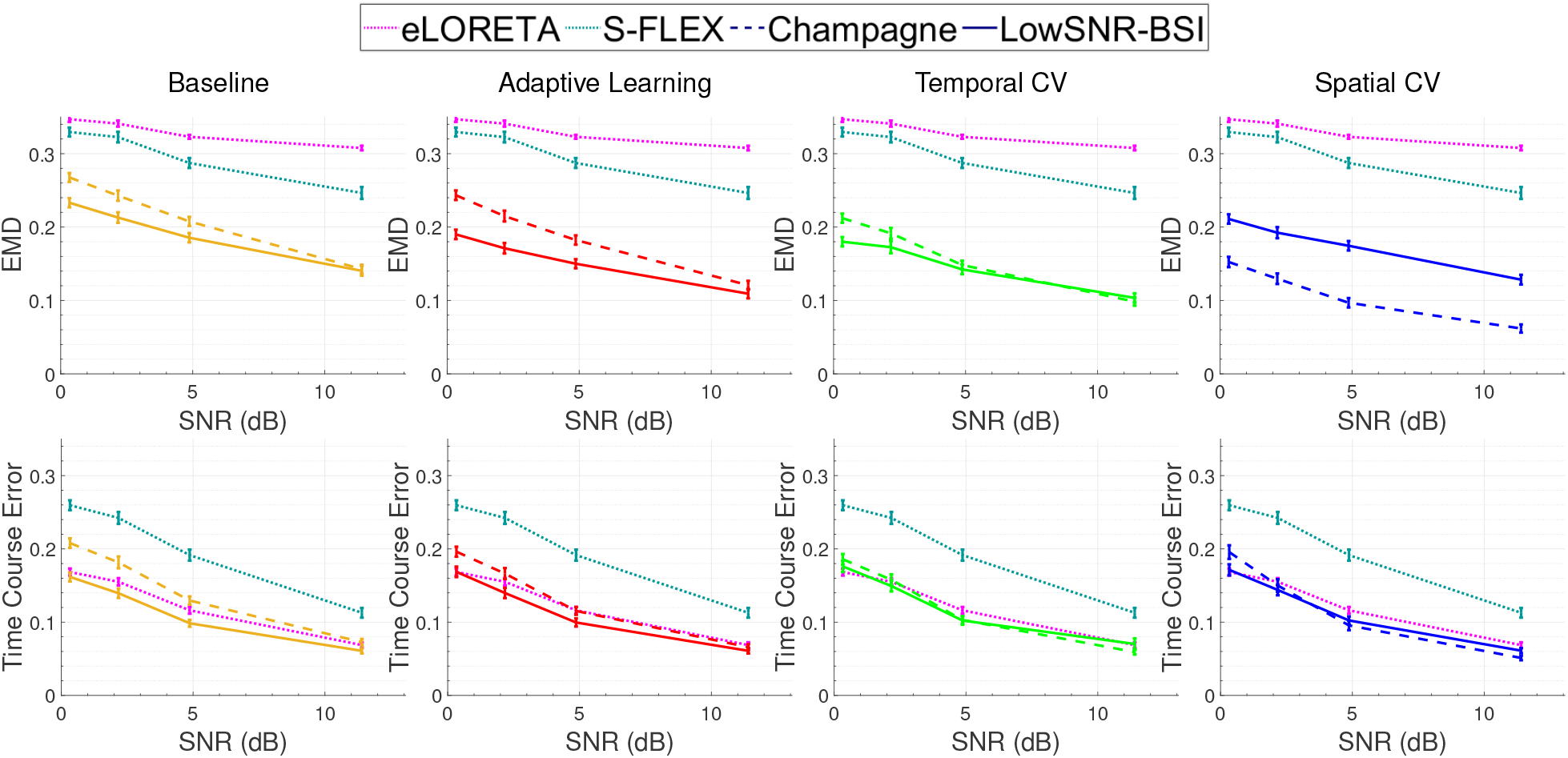
Source reconstruction performance of Champagne (dashed line) and LowSNR-BSI (solid line) for four SNR values (SNR = {0.33, 2.17, 4.87, 11.40} dB). The noise variance was estimated from baseline data as well as using adaptive learning, spatial and temporal CV. Spatial reconstruction error was measured in terms of the earth-mover’s distance and is shown in the upper row, while time course reconstruction error is shown in the lower row.

Note that the LowSNR-BSI surrogate function in the baseline mode can provide a tight upper-bound for the original non-convex function only when SNR is equal to zero. For non-zero SNR, the current theory unfortunately does not apply; however, it is clear from our empirical results that the LowSNR-BSI surrogate function remains advantageous also in non-zero low-SNR regimes. As Figure 3 demonstrates, we observe performance improvements of LowSNR-BSI over Champagne in the baseline mode for SNRs up to around 8 dB. After this point, the surrogate functions of LowSNR-BSI and Champagne both appear to be able to approximate the non-convex loss with a similar degree of precision; thus, their performance with respect to the evaluation metrics overlap as can be seen in the right column of Figure 1.

It can further be observed that S-FLEX yields higher spatial localization accuracy (lower EMD) than eLORETA, while eLORETA yields higher temporal accuracy (lower time course error) than S-FLEX across all SNR values. With respect to spatial accuracy, both approaches, however, are consistently outperformed by Champagne and LowSNR-BSI. Note that the superior spatial reconstruction of sparsity-inducing algorithms (Champagne, LowSNR-BSI and S-FLEX) compared to eLORETA is expected here, because the simulated spatial distributions are indeed sparse. The superiority of SBL methods (Champagne, LowSNR-BSI) over S-FLEX that is observed here confirms observations and theoretical considerations made in [8], [27], [28]. eLORETA shows comparable temporal reconstruction performance as LowSNR-BSI and Champagne, while S-FLEX is outperformed by all other methods.

The convergence behavior of the different SBL variants discussed and introduced in Sections III–V is illustrated in Figure 4. LowSNR-BSI variants have faster convergence rates at the early stage of the optimization procedure compared to standard Champagne as well as Champagne with MacKay updates. They, however, reach lower negative log-likelihood values eventually, which indicates that they find better maxima of the model evidence. Furthermore, the adaptive-learning variants of Champagne and LowSNR-BSI reach lower negative log-likelihood values than their counterparts estimating the noise variance from baseline data, suggesting that learning the noise variance, or in other words overestimating the noise variance, improves the reconstruction performance through better model evidence maximization.

**Fig. 4.**
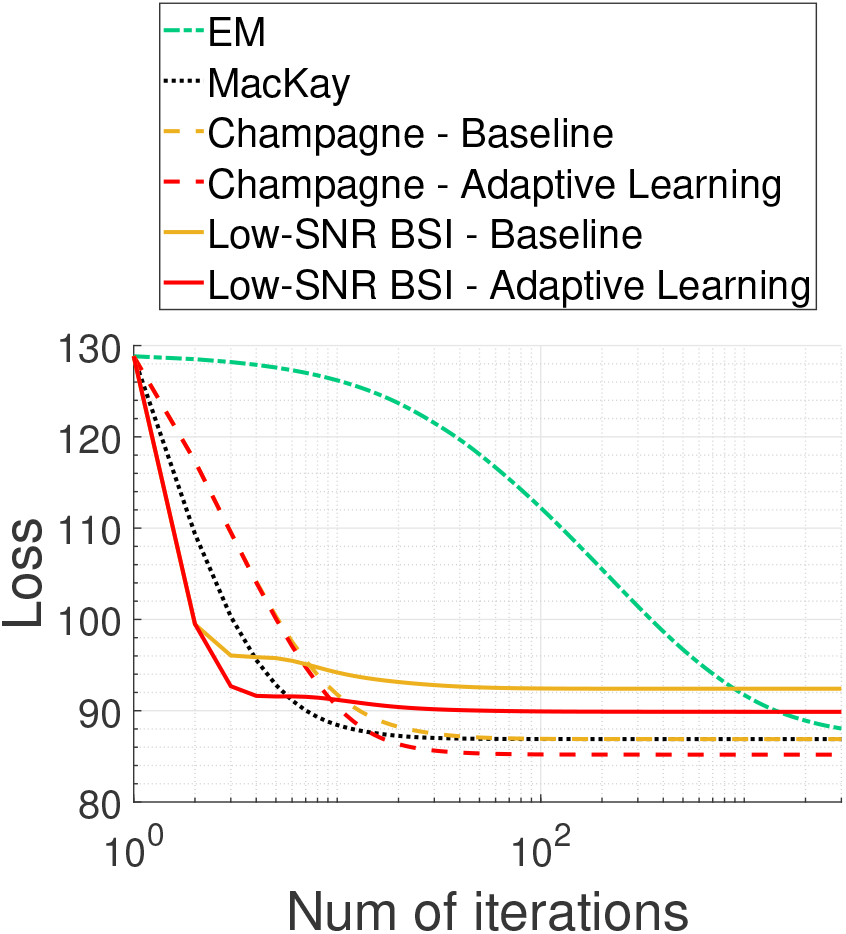
Convergence behavior of LowSNR-BSI as well as Champagne using the standard (convex-bounding based) updates (Champagne) as well as EM and MacKay updates. For standard Champagne and LowSNR-BSI, the use of a fixed noise variance estimated from baseline data is compared with adaptive noise learning. LowSNR-BSI variants have faster convergence rate at early stages of the optimization procedure, but later converge to less optimal log-likelihood values. Adaptive learning variants of Champagne and LowSNR-BSI reach better log-likelihood values than their counterparts using a fixed noise variance estimated from baseline data.

Note that the plots in Figure 4 demonstrate the convergence behaviour of MM algorithms for only one single experiment. We conducted another experiment (see Appendix I), in which the simulation was carried out 100 times using different instances of source distributions and initializations. The final negative log-likelihood loss – attained after convergence – and runtimes of all methods were calculated. The median and inter-quartile ranges over 100 randomized experiments of these performance metrics are reported in Figure 8, which confirms the observations made here.

## VII. Analysis of Auditory Evoked Fields (AEF)

The MEG data used here were acquired in the Biomagnetic Imaging Laboratory at the University of California San Francisco (UCSF) with a CTF Omega 2000 whole-head MEG system from VSM MedTech (Coquitlam, BC, Canada) with 1200 Hz sampling rate. The neural responses of one subject to an Auditory Evoked Fields (AEF) stimulus were localized. The AEF response was elicited with single 600 ms duration tones (1 kHz) presented binaurally. The data were averaged across 120 trials (after the trials were time-aligned to the stimulus). The pre-stimulus window was selected to be −100 ms to 5 ms and the post-stimulus time window was selected to be 5 ms to 250 ms, where 0 ms is the onset of the tone. Further details on this dataset can be found in [27], [28], [84]. The lead field for each subject was calculated with NUTMEG (http://bil.ucsf.edu) using a single-sphere head model (two spherical orientation lead fields) and an 8 mm voxel grid.

The results presented in Section VI have been obtained for the scalar setting, where the orientation of the brain sources are assumed to be perpendicular to the surface of cortex and, hence, only the scalar deflection of each source along the fixed orientation needs to be estimated. In real data, surface normals are hard to estimate or even undefined in case of volumetric reconstructions. Consequently, we model each source here as a full 3-dimensional current vector. This is achieved by introducing three variance parameters for each source within the source covariance matrix, 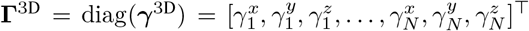. As all algorithms considered here model the source covariance matrix **Γ** to be diagonal, this extension can be readily implemented. Correspondingly, a full 3D leadfield matrix, **L**^3D^ ∈ ℝ^*M* ×3*N*^, is used.

Figure 5 shows the reconstructed sources of the AEF of one subject using conventional Champagne with pre-estimated 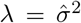, adaptive noise learning, and spatial CV. LowSNR-BSI with pre-estimated 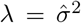 was also included in the comparison. Shown in the top panel are the reconstructions at the time of the maximal deflection of the auditory N100 component (shown in bottom panel).

**Fig. 5.**
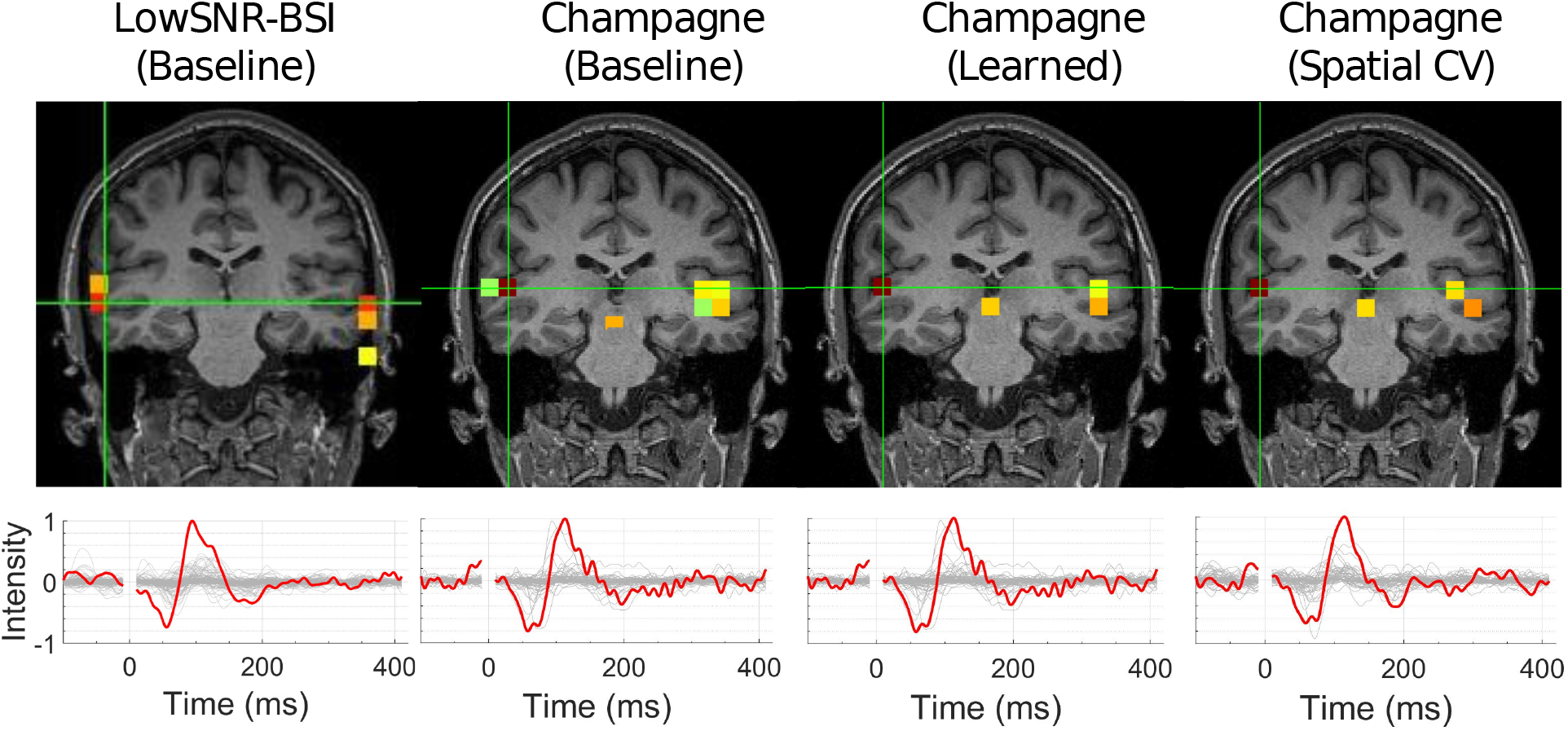
Analysis of auditory evoked fields (AEF) of one subject using conventional Champagne with pre-estimated 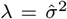, adaptive noise learning, and spatial CV as well as LowSNR-BSI. Shown in the top panel are the reconstructions at the time of the maximal deflection of the auditory N100 component (shown in bottom panel). All reconstructions show sources at the expected locations in the left and right auditory cortex.

All reconstructions are able to correctly localize bilateral auditory activity to Heschel’s gyrus, which is the location of the primary auditory cortex. Note that an additional source in the midbrain, which is indicated by all three Champagne variants, is absent for LowSNR-BSI.

We tested the reconstruction performance of all methods for random subsets of 10, 20, 40, 60, and 100 trials. As Figure 6 shows, the proposed noise learning variants of Champagne as well as LowSNR-BSI can correctly localize bilateral auditory activity to Heschl’s gyrus even when using as few as 10 trials. Focusing on the low-SNR regime, Figure 7 shows seven reconstructions for random selections of 10 trials. LowSNR-BSI as well as all proposed noise learning variants of Champagne consistently show sources at the expected locations in the left and auditory cortices, where both cortices are jointly identified in the majority of experiments.

**Fig. 6.**
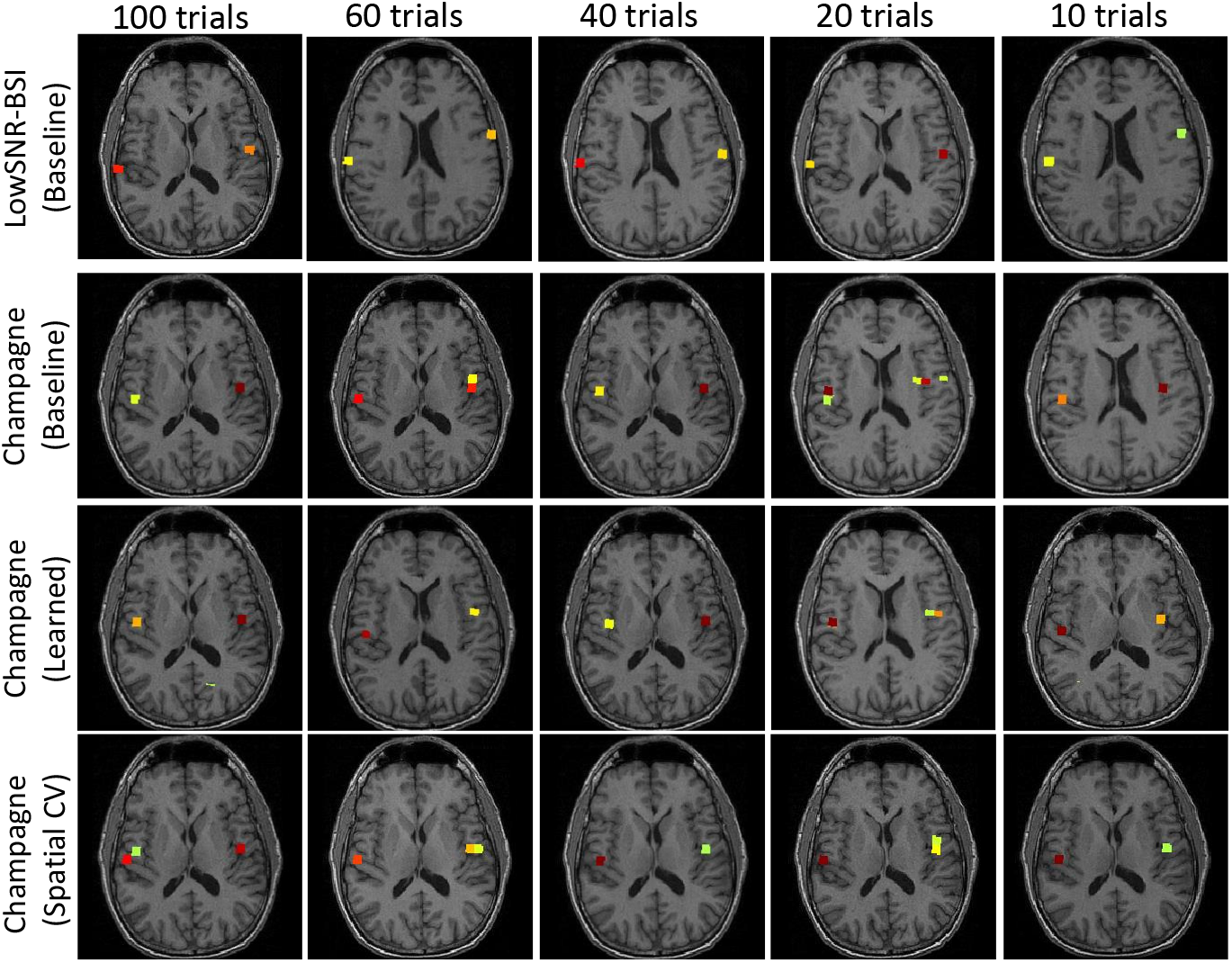
Analysis of auditory evoked fields (AEF) of one subject using conventional Champagne with pre-estimated 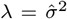, adaptive noise learning, and spatial CV as well as LowSNR-BSI, tested with the number of trials limited to 10, 20, 40, 60, and 100. All proposed noise learning reconstructions show sources at the expected locations in the left and right auditory cortices.

**Fig. 7.**
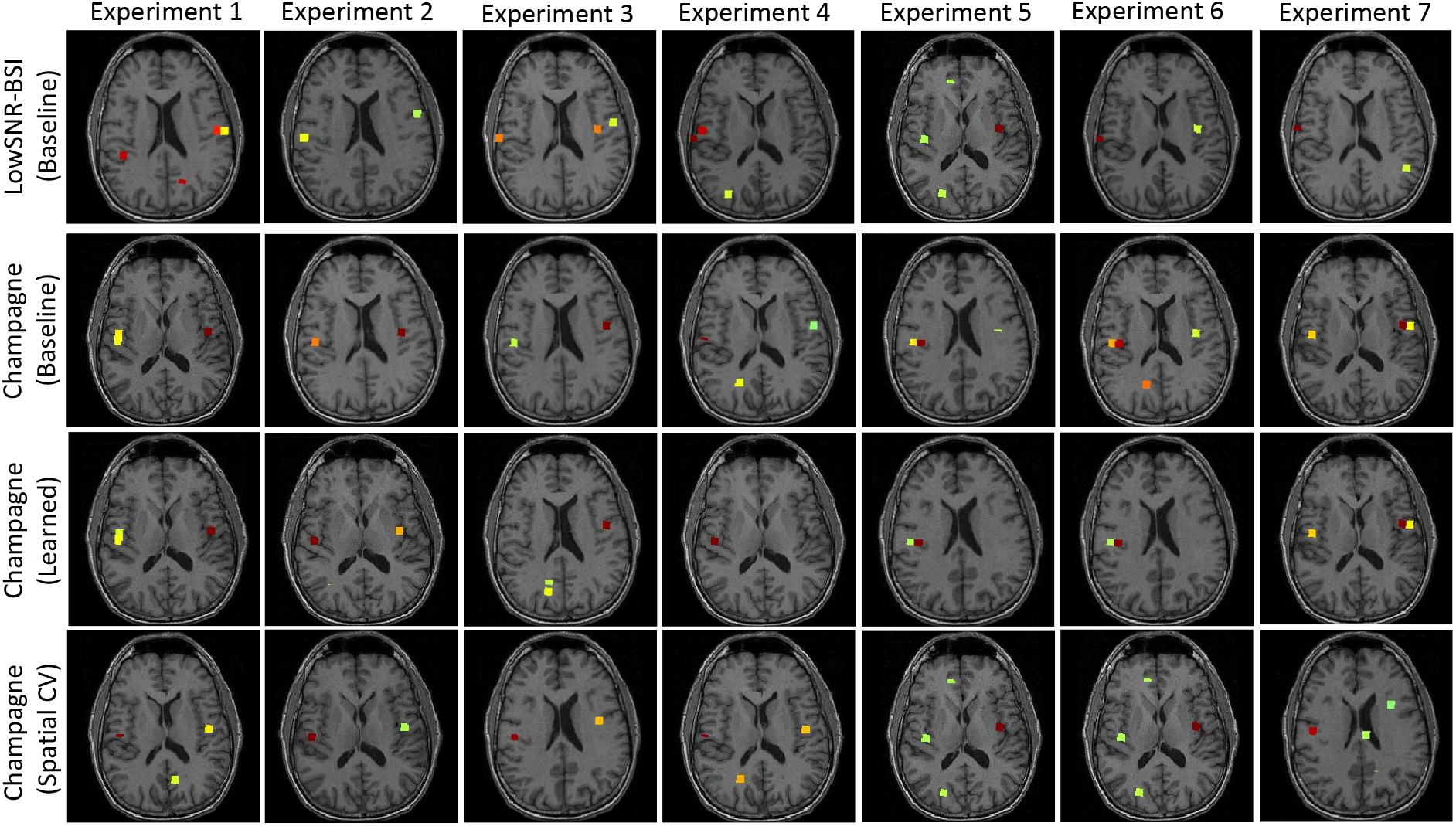
Analysis of auditory evoked fields (AEF) of one subject using conventional Champagne with pre-estimated 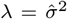, adaptive noise learning, and spatial CV as well as LowSNR-BSI, tested with the number of trials limited to 10. Each column shows an experiment with a random selection of 10 trials. LowSNR-BSI as well as all proposed noise learning variants of Champagne always show sources at the expected locations in the left or right auditory cortex. In the majority of experiments, both cortices are jointly identified.

**Fig. 8.**
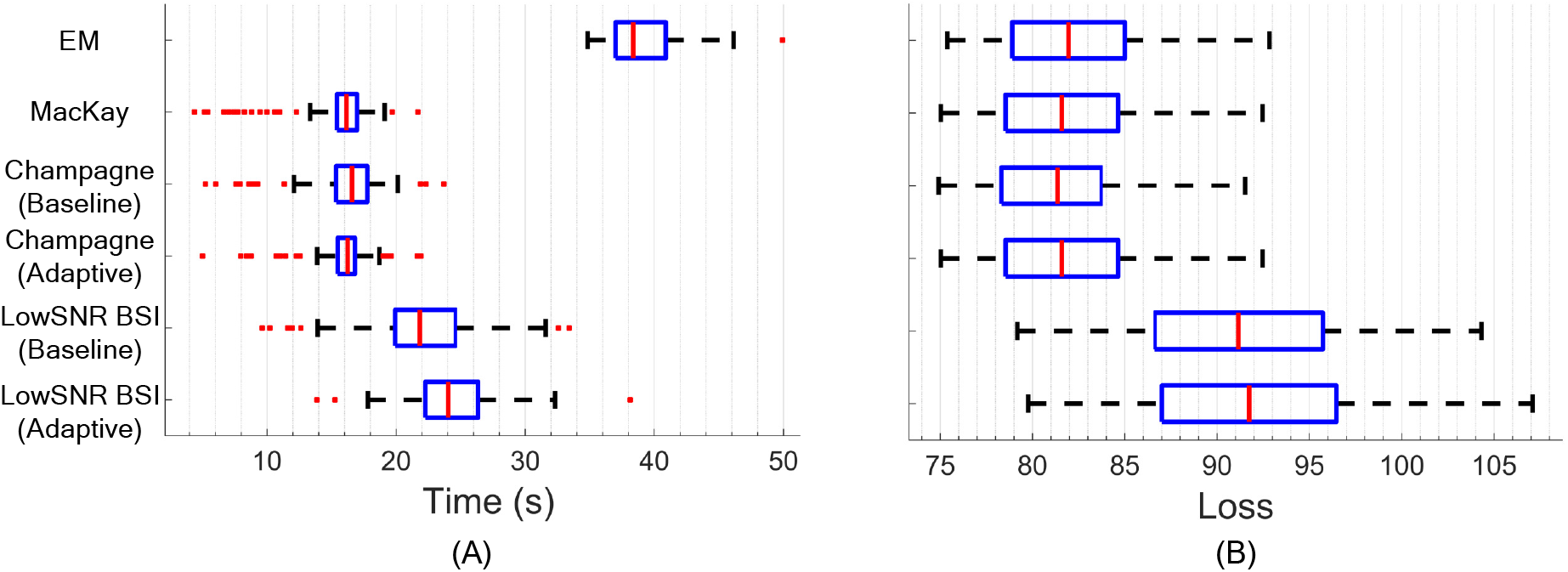
(A) Runtime and (B) Type-II negative log marginal likelihood loss attained after convergence of different variants of LowSNR-BSI and Champagne, as well as using EM and MacKay updates. Shown are the median and inter-quartile ranges over 100 randomized experiments.

## VIII. Discussion

We have provided a unifying theoretical platform for deriving different sparse Bayesian learning algorithms for electromagnetic brain imaging using the Majorization-Minimization (MM) framework. First, we demonstrated that the choice of upper bounds of the Type-II non-convex loss function within the MM framework influences the reconstruction performance and convergence rates of the resulting algorithms. Second, focusing on commonly occurring low-SNR settings, we derived a novel Type-II Bayesian algorithm, LowSNR-BSI, using a novel convex bounding MM function that converges to the original loss function as the SNR goes to zero. We demonstrated the advantage of LowSNR-BSI over existing benchmark algorithms including Champagne, eLORETA and S-FLEX. Consistent with the theoretical considerations, the advantage of LowSNR-BSI over Champagne decreases with increasing SNR. Third, we have derived an analytic solution that allows us to estimate the noise variance jointly within the source estimation procedure on the same (task-related) data that are used for the reconstruction. We have also adopted cross-validation schemes to empirically estimate the noise variance from hold-out data through a line search. We have proposed spatial and temporal CV schemes, where either subsets of EEG/MEG channels or recorded samples are left out of the source reconstruction, and where the noise variance is selected as the minimizer of a divergence between model and hold-out data. We also demonstrate that precise knowledge of the noise variance is required in order to determine the optimal algorithm performance. Finally, according to our empirical results, all three proposed techniques for estimating the noise variance lead to superior source reconstruction performance compared to the setting in which the noise variance is estimated from baseline data.

### A. Cross-validation vs. adaptive noise learning

Spatial CV for Champagne and Temporal CV for LowSNR-BSI achieved the best performances and are generally applicable to any distributed inverse solution. Their long computation time can, however, be challenging as their computational complexity is drastically higher (around two orders of magnitude) than using baseline data or adaptive learning schemes. The high complexity of CV techniques is a potential limitation in settings where the efficiency of the algorithm or immediate access to the outcome is crucial. What is more, this approach quickly becomes infeasible if more than one parameter needs to be estimated through a grid search. In contrast, the computational complexity of the proposed noise level estimation scheme using adaptive learning is of the same order as the complexity of the baseline approach. Moreover, we have successfully extended this approach to the estimation of heteroscedastic noise, where a distinct variance is estimated for each M/EEG sensor [28]. Hence, the adaptive-learning approach can be seen as an advancement of the baseline algorithm that combines performance improvement and computational efficiency. It is also worth noting that the computational complexity of CV techniques heavily relies on tunable parameters such as the number of folds/splits of the data and the total number of candidate points in the grid search.

### B. Interpretation of Type-I and Type-II loss functions as divergences

We have pointed out (see Section V-B and Appendix A) that Type-I and Type-II Bayesian approaches implicitly use different metrics to compare the empirical sensor-space observations to the signal proportion explained by the reconstructed brain sources. Type-I approaches measure first-order differences between modeled and reconstructed time series using variants of the MSE, while Type-II approaches amount to using the log-det Bregman divergence to measure differences in the second-order statistics of the empirically observed and modeled data as summarized in the respective covariance matrices. While the connection between the Type-II loss function and the log-det Bregman divergence has been investigated and exploited in numerous forms such as *Stein’s loss* [50] or the *graphical Lasso* [78], [85], [86], and has found applications in disciplines such as information theory and metric learning [87], [88], wireless communication [21], and signal processing [26], [89], [90], it has not received much attention in the BSI literature to the best of authors’ knowledge. Here, we have used this insight to devise a novel cross-validation scheme, temporal CV, in which model fit is measured in terms of the log-det Bregman divergence (or, Type-II likelihood) on held-out samples. In contrast, the previously introduced spatial CV uses the mean-squared error to measure out-of-sample model fit. Importantly, however, this difference does not imply that the application of spatial CV is restricted to Type-I approaches or that the use of temporal CV is restricted to Type-II approaches. Rather, both approaches are universally applicable. In fact, it is straightforward to evaluate the Type-I likelihood based on the source times series reconstructed with Type-II methods. Conversely, it is also possible to estimate the Type-II likelihood for Type-I approaches such as S-FLEX. Here, the model source and noise covariances are first estimated from the reconstructed sources as 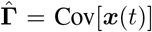 and 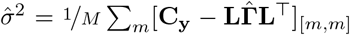, after which **Σ_*y*_** can be calculated. The optimal Type-I regularization parameter is then selected as the minimizer of 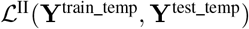 in Eq. (31).

### C. Limitations and future work

One limiting assumption of the current work is that the activity of the sources is modeled to be independent across voxels, spatial orientations, and time samples. Analogously, the noise is assumed to be independent across times samples, and homoscedastic (independent with equal variance across sensors). These assumptions merely act as prior information whose purpose is to bias the inverse reconstruction towards solutions with lower complexity. Thus, they do not prevent the reconstruction of brain and noise sources with more complex structure if the observed data are inconsistent with these priors. On the other hand, modeling dependency structures that are in fact present in real data has the potential to substantially improve the source reconstruction. We have recently proposed adaptive noise learning algorithms that relax the rather unrealistic assumption of homoscedastic noise [28]. Going further, it would be possible to also model spatial covariances of the sources between voxels and/or between source orientation within voxels, which would encode the realistic assumption that individual brain regions do not work in isolation. Similary, the spatial covariance structure of the noise could be modeled in order to accommodate spatially distributed artifacts due to, for example, heart beat or line noise interference. Finally, electrophysiological data are known to possess a complex intrinsic autocorrelation structure, which is not modeled by the majority of existing BSI algorithms. We have recently proposed ways to also learn temporal correlations within the Type-II framework and have obtained promising results with respect to time course reconstruction [24], [91].

## IX. Conclusion

We have provided a unifying theoretical platform for deriving different sparse Bayesian learning algorithms for electromagnetic brain imaging using the Majorization-Minimization (MM) framework. This unification perspective not only provides a useful theoretical framework for comparing different algorithms in terms of their convergence behavior, but also provides a principled recipe for constructing novel algorithms with specific properties by designing appropriate bounds of the Bayesian marginal likelihood function. Building on MM principles, we then proposed a novel method called *LowSNR-BSI* that achieves favorable source reconstruction performance in low signal-to-noise-ratio settings. Recognizing the importance of noise estimation for algorithm performance, we present both analytical and cross-validation approaches for noise estimation. Empirically, we show that the monotonous convergence behavior predicted from MM theory is confirmed in numerical experiments. Using simulations, we further demonstrate the advantage of LowSNR-BSI over conventional Champagne in low-SNR regimes, and the advantage of learned noise levels over estimates derived from baseline data. To demonstrate the usefulness of our novel approach, we show neurophysiologically plausible source reconstructions on averaged auditory evoked potential data.

Our characterization of the Type-II likelihood as a divergence measure provides a novel perspective on the construction of BSI algorithms and might open new avenues of research in this field. It is conceivable that alternative divergence metrics can be used for solving the M/EEG source reconstruction problem in the future by modeling specific neurophysiologically valid aspects of similarity between data and model output. Promising metrics in that respect are information divergences such as Kullback-Leibler (KL) [92], Rényi [90], Itakura-Saito (IS) [93] and *β* divergences [94]–[97] as well as transportation metrics such as the Wasserstein distance between empirical and statistical covariances (e.g., [98]–[101]).

Although this paper focuses on electromagnetic brain source imaging, Type-II methods have also been successfully developed in other fields such as direction of arrival (DoA) and channel estimation in wireless communications [18], [19], [21], [102], Internet of Things (IoT) [103], [104], robust portfolio optimization in finance [22], covariance matching and estimation [105]–[112], graph learning [113], and brain functional imaging [92]. The methods introduced in this work may also prove useful in these domains.

## Appendix

## A. Bregman Divergence Formulation of the Type-II Loss Function

We start by recalling the definition of log-det Bregman matrix divergence - also known as Stein’s loss [50] - between any two *M* × *M* positive semidefinite (PSD) matrices **Q** and **W**:

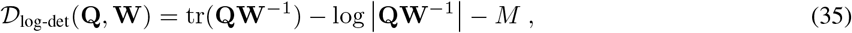

where the “log-det” Bregman matrix divergence in (35) is an special case of Bregman matrix divergence [79], where −log |·| is selected as a strictly convex function. By substituting **C_y_** and **Σ_y_** in (35) instead of **Q** and **W**, the *log-det* Bregman matrix divergence can be written as follows [21], [26], [78], [85]–[88], [114], [115]:

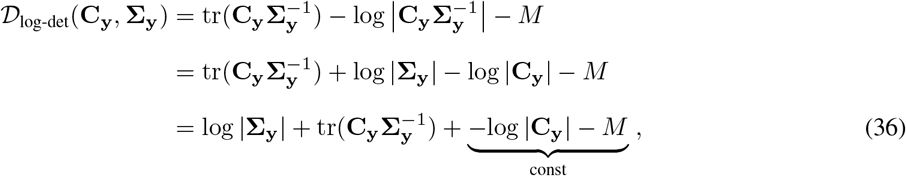

where (36) is the same as (13) up to a constant. Note that log |**C_y_**| does not depend on ***γ*** and is, therefore, treated as a constant value here.

## B. Proof of Corollary 1

*Proof.* To verify the descending trend in the MM framework, it is sufficient to show that *f* (**u**^*k*+1^) ≤ *f* (**u**^*k*^). To this end, we have *f* (**u**^*k*+1^) ≤ *g*(**u**^*k*+1^|**u**^*k*^) from condition [A2]. Condition [A3] further states that *g*(**u**^*k*+1^|**u**^*k*^) ≤ *g*(**u**^*k*^|**u**^*k*^), while *g*(**u**^*k*^|**u**^*k*^) = *f* (**u**^*k*^) holds according to [A1]. Putting everything together, we have:

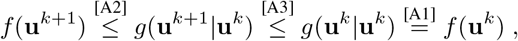

which concludes the proof.

## C. Proof of Proposition 1

*Proof.* We first show that the objective function of the M-step is derived by upper-bounding the negative log-likelihood, − log *p*(**Y**|***γ***), using Jensen’s inequality (J):

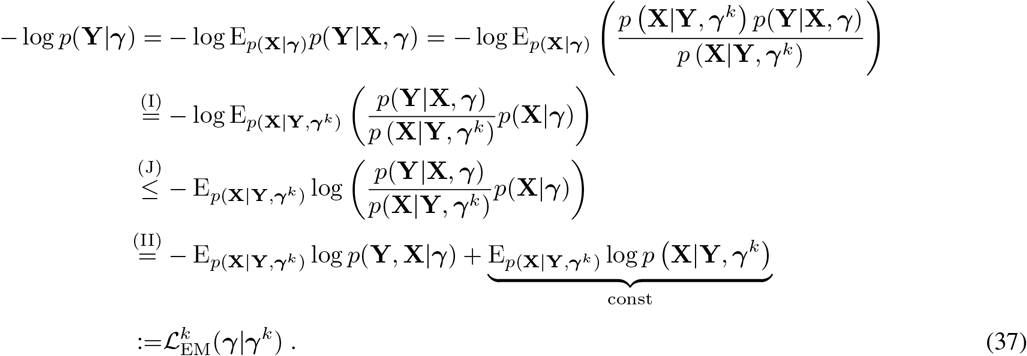

The resulting bound is a majorizing function for − log *p*(**Y**|***γ***), so that condition [A2] holds. Note that the term 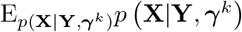 does not depend on ***γ*** and, therefore, does not influence the optimization. According to the definition of Jensen’s inequality, the equality constraint – condition [A1] – holds if and only if the argument of the convex function is a constant. Therefore, to establish the equivalence of both sides of (J) when *γ* = *γ^k^*, it is sufficient to show that the argument of the log function, 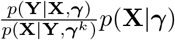, is constant when *γ* = *γ^k^*. This can be verified by invoking Bayes rule:

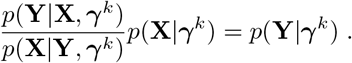

Since *p*(**Y**|***γ***^*k*^) is a constant, equality condition [A1] holds.

After inserting the analytic form of − log *p*(**Y**, **X**|***γ***) in Eq. (24):

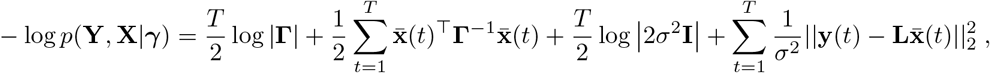

we are ready to prove that 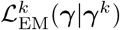 fulfills condition [A3]. We have:

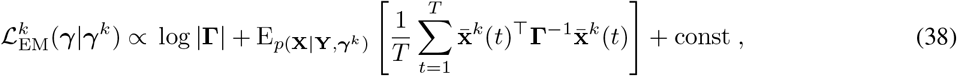

where const comprises all terms of Eq. (25) that are not a function of ***γ***. To prove that 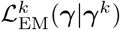 satisfies condition [A3], we need to show that 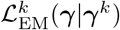 reaches to its global minimum in each MM iteration. This can be easily guaranteed if Eq. (38) is convex. While the second term in (38) is convex, the first term, log |**Γ**|, is in fact concave, which hampers conclusions concerning the convexity of their sum. However, we can use the concept of *geodesic convexity* or *g-convexity* from non-Euclidean and geometric optimization, which enables us to prove that any local minimum of Eq. (38) is actually a global minimum. For the sake of brevity, we will omit a detailed theoretical introduction of g-convexity, and only borrow the following required propositions, Propositions 5 and 6, from the literature (an interested reader can refer to [89, Chapter 1] for a gentle introduction to this topic, and to [116, Chapter 2] [117]–[123] for more in-depth technical details). Now, we state the following preliminary results:

### Proposition 5.

*The function* log |**Γ**| *is g-convex in* **Γ**, *where* **Γ***belongs to the manifold of positive definite (PD) matrices.*

*Proof.* A detailed proof can be found in [89, Lemma. 1.13]. The main idea is to leverage the geodesic **Q**_*q*_ = **VD**^*q*^**V**^⊤^, *q* ∈ [0, 1] between two matrices, **Q**_0_ = **VV**^⊤^ and **Q**_1_ = **VDV**^⊤^, in order to transfer the problem into the following form:

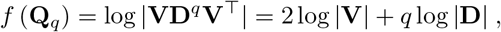

where *f* (**Q**_*q*_) is a linear function and, therefore, convex in *q*.

### Remark 7.

*The log-determinant function is concave in classical Euclidean analysis. However, Proposition 5 demonstrates that it is g-convex with respect to the PD manifold.*

### Proposition 6.

*Any local minimum of a g-convex function over a g-convex set is a global minimum. Proof.* A detailed proof is presented in [117, Theorem 2.1].

Given that g-convexity is an extension of classical convexity to non-Euclidean geometry, it is straightforward to show that all convex functions are also g-convex, where the geodesics between pairs of matrices are simply line segments. Therefore, given Proposition 5, we can conclude that Eq. (38) is g-convex; hence, any local minimum of 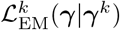 is a global minimum according to Proposition 6. This proves that condition [A3] is fulfilled and completes the proof of Proposition 1.

## D. Proof of Proposition 2

*Proof.* We start by recalling 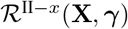 in Eq. (14):

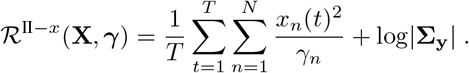

Based on [16, Example 2], [A2] can be directly inferred from the concavity of the log-determinant function and its first-order Taylor expansion around the value from the previous iteration, 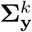, which leads to the following inequality:

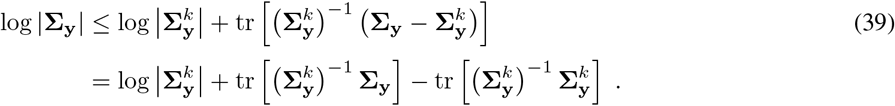

Note that the first and last term in (39) do not depend on ***γ***; hence, they can be ignored in the optimization procedure. Conditions [A1] and [A4] are automatically satisfied by construction because the majorizing function is obtained through a Taylor expansion around 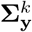. Concretely, [A1] is satisfied because the equality in Eq. (39) holds for 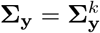. Similarly, [A4] is satisfied because the gradient of log |**Σ_y_**| at point 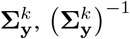, defines the linear Taylor approximation 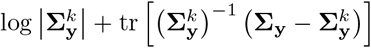. Thus, both gradients coincide in 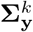 by construction. Now, we show that [A3] can be satisfied easily using standard optimization algorithms by proving that 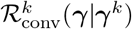 is a convex function with respect to ***γ***. To this end, we rewrite Eq. (26):

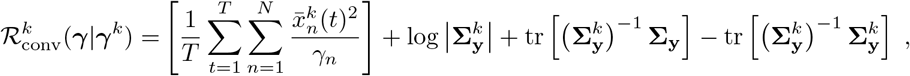

as follows:

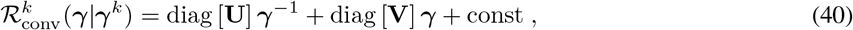

where 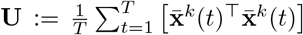 and 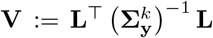 are defined as parameters that do not depend on ***γ***. The term const also collects constant terms in (39), i.e. 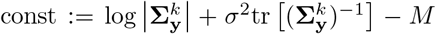. Besides, 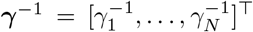 is defined as the element-wise inversion of ***γ***. The convexity of 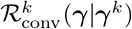 can be directly inferred from the convexity of diag [**U**] ***γ***^−1^ and diag [**V**] ***γ*** with respect to ***γ*** [124, Chapter. 3]. The convexity of 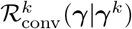, which ensures that condition [A3] can be satisfied using standard optimization, along with fulfillment of conditions [A1], [A2] and [A4], ensure that Theorem 1 holds.

In order to establish the equivalence of the MM algorithm using the majorization function Eq. (26) and the convex-bounding based Champagne variant presented in Section II-C2, we here decompose **Σ_y_** into rank-one matrices as introduced in [125]. The first term of Eq. (26) can be reformulated as follows:

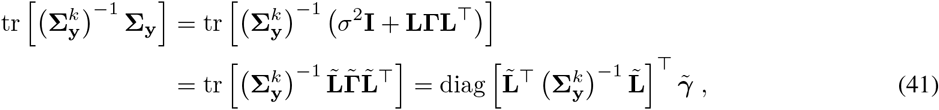

where 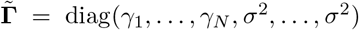, and 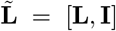. Since we are optimizing Eq. (26) with respect to *γ_n_*, for *n* = 1, …, *N*, the elements of 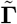 and 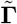 related to the sensor noise *σ*^2^ vanish. Thus, by inserting Eq. (41) into Eq. (26), taking the derivative with respect to *γ_n_*, for *n* = 1, …, *N*, and setting it to zero,

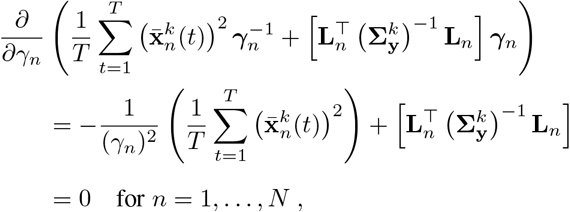

where **L**_*n*_ denotes the *n*-th column of the lead field matrix, we obtain an update rule in terms of the original variables **Γ** and **L**:

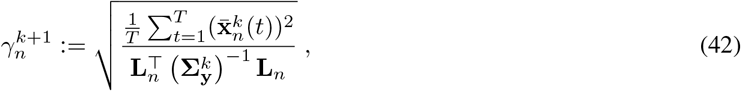

which is identical to the update rule of the convex-bounding based approach discussed in Section II-C2, Eqs. (17)– (19).

## E. Proof of Proposition 3

*Proof.* The proof that conditions [A1]–[A4] are satisfied is directly analogous to that of Proposition 2; therefore, it is omitted here. The equivalence of the Champagne variant based on MacKay updates [12, Section III.A-2] presented in Section II-C3 and the solution derived within the MM framework can be derived by transforming the update rule Eq. (42) into a fixed-point iteration of the form ***γ***^*k*+1^ = *f* (***γ***^*k*^), which is an alternative way of minimizing the same surrogate function (Eq. (26)). By squaring the left and right hand sides of Eq. (42), one can divide both sides by 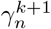 and re-interpret the term on the right hand side as the estimate from the previous (*k*-th) iteration:

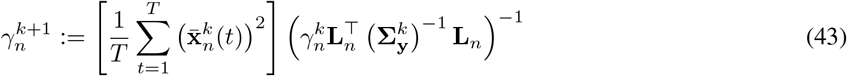

for *n* = 1, …, *N*. This is indeed identical to the MacKay update in Eq. (22), which concludes the proof.

## F. Proof of Proposition 4

*Proof.* (following [19, Appendix C-A]) Without loss of generality, we here consider the case *σ*^2^ = 1, which can be obtained by normalizing the sensor and source covariance matrices by 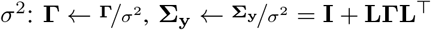. Also, due to the concavity of the log(·) function and by using a Taylor expansion around point *a*, we have:

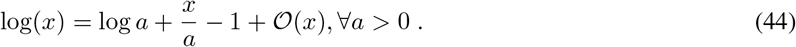

Assuming that **LΓL**^⊤^ has an eigenvalue decomposition **LΓL**^⊤^ = **UPU**^⊤^ with **P** = diag(*p*_1_, …, *p_M_*), the majorizing function 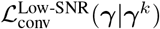 as well as Eq. (28) are derived as follows:

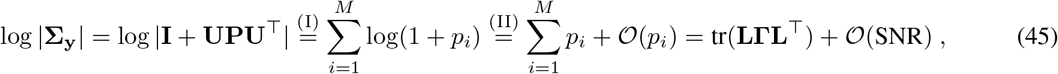

where the *p_i_*, for *i* = 1, …, *M* denote the diagonal elements of **P**, which are equivalent to the eigenvalues of **LΓL**^⊤^. The term 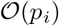 represents the second and higher-order residuals of the Taylor expansion. Note that (45)-(I) is obtained by expanding **P** over its diagonal elements, while (45)-(II) is derived by exploiting the concavity of the log(.) function and its first-order Taylor expansion around *a* = 1 based on Eq. (44). Given the eigenvalue decomposition of **LΓL**^⊤^ = **UPU**^⊤^ and the normalization with respect to the noise variance, the sum over all eigenvalues of **LΓL**^⊤^, i.e., 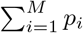, represents the ratio between the power of the signal and the power of the noise; hence, one can replace 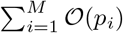 in Eq. (45) with 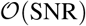. To elaborate this more, note that given *σ*^2^ = 1, we have 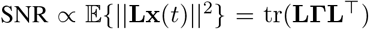, where 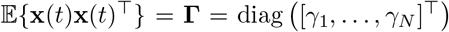 due to the independence between voxels. Therefore, 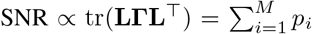, as the sum of the eigenvalues of a matrix is equal to its trace.

As we have shown that 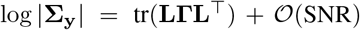, condition [A2] holds and 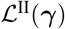 converges to 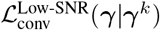 when SNR → 0. Moreover, as Eq. (28) is constructed using a linear Taylor approximation, [A1] and [A4] hold due to the same arguments made in the proof of Proposition 2. It remains to be shown that condition [A3] can be easily fulfilled due to the convexity of 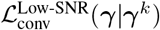. To this end, we exploit the following key relationship between the sensor and source space covariances:

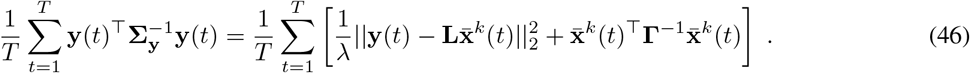

By replacing 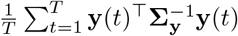 in Eq. (27) with its source space equivalence in (46), we have:

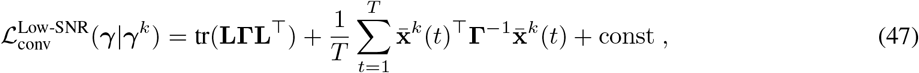

where const denotes the terms that do not depend on ***γ***. Reformulating (47) as

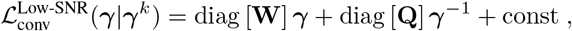

with 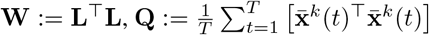 and 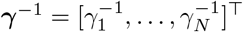 proves the convexity of 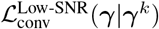 using the same arguments made for proving convexity in Proposition 2. Thus, we have shown that conditions [A1]–[A4] hold, which concludes the proof.

## G. Detailed Derivation of the LowSNR-BSI Algorithm

To find the optimal value of ***γ*** = [*γ*_1_, …, *γ_N_*]^⊤^, we take the derivative of 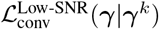 in (27) with respect to each *γ_n_* for *n* = 1, …, *N* :

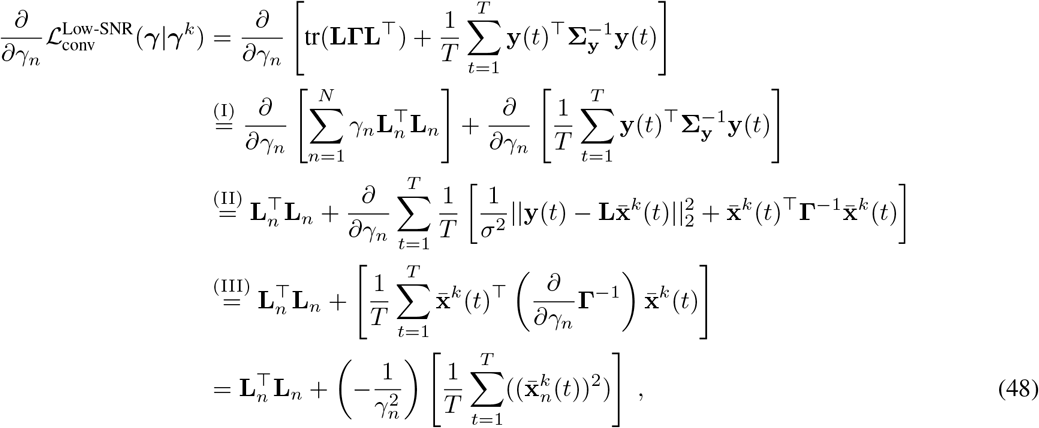

where Eq. (48)-I is derived based on a *sum-of-rank-one matrices* reformulation of the term tr(**LΓL**^⊤^) by exploiting the diagonal structure of **Γ**. Equality (48)-II is the direct implication of the duality between ***γ***-space and **X**-space that has been pointed out in (14). Finally, 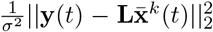 does not appear in (48)-III and is ignored since it does not depend on ***γ***. Setting the derivative in Eq. (48) to zero yields the following closed-form update for ***γ*** = [*γ*_1_, …, *γ_N_*]^⊤^:

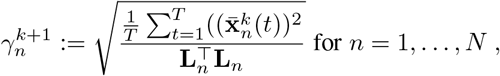

which is identical to the update rule in Eq. (29). This completes the derivation of the LowSNR-BSI algorithm.

## H. Proof of Theorem 2

*Proof.* We start by taking the derivative of 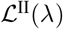 with respect to *λ*:

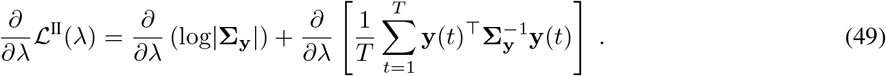

We first calculate the first term, 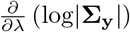. Using the matrix inversion equality

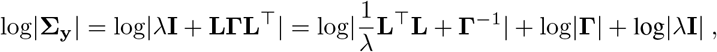

we have

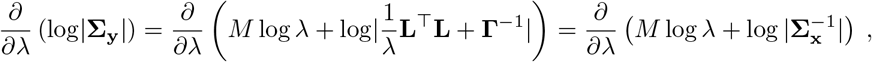

where the term log|**Γ**| is omitted since it is does not depend on *λ*. Then, the derivative of log|**Σ_y_**| with respect to *λ* can be obtained as follows:

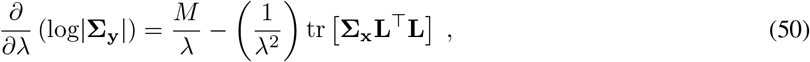

where the second term in (50) is derived according to the equality 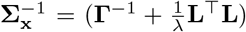, which holds for the inverse of the posterior covariance in Eq. (10) [49, Chapter 4]:

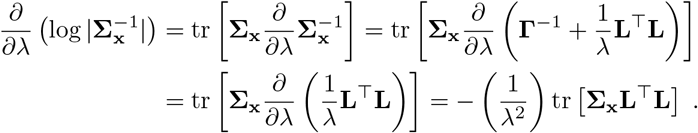

In the next step, we calculate the derivative of the second term in Eq. (49) using the following key relation between the sensor and source space covariances presented in Appendix F. Given (46), we have

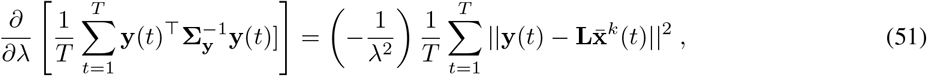

where the term 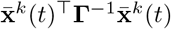 is neglected since it does not depend on *λ*. Let **Γ**^*k*^ and 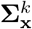 be fixed values obtained in the (*k*)-th iteration. Then, by substituting Eqs. (50) and (51) into Eq. (49), we have:

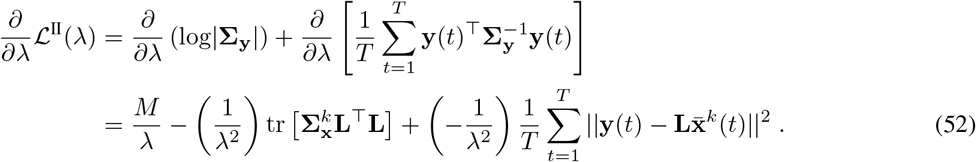

By expressing 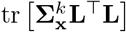 in terms of the values at the (*k*)-th iteration according to the following matrix equality [71]:

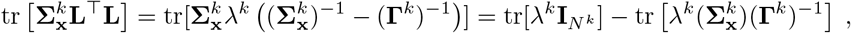

Eq. (52) can be reformulated as follows:

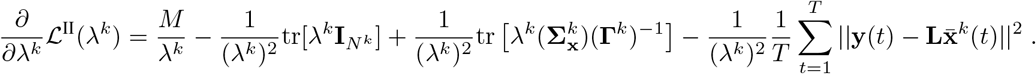

Note that *N^k^* denotes the number of non-zero voxels at the (*k*)-th iteration. Now by setting the derivative to zero, the update rule for *λ* at the (*k* + 1)-th iteration is obtained as

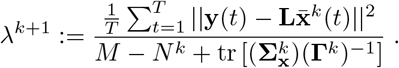

This completes the proof.

## I. A Statistical Analysis of Computational Complexity and Convergence Behaviour of MM Methods

Here, we conducted an experiment, in which the simulation presented in Figure 4 was carried out 100 times using different instances of source distributions and initializations. The final negative log-likelihood loss – attained after convergence – and runtimes of all methods were calculated. The median and inter-quartile ranges over 100 randomized experiments of these performance metrics are reported in Figure 8.

As demonstrated in Figure 4, the EM algorithm indeed needs a larger number of iterations for convergence than its peer MM variants, which eventually results in longer runtimes and higher computational complexity if we measure runtime in units of seconds, demonstrated in Figure 8-(A). The overall computation complexity in each iteration of the EM, however, is comparable to the other MM variants. Even though an additional operation for calculating the posterior matrix of the sources, **Σ_x_**, is involved in each iteration of the EM algorithm – which operates in the high-dimensional source space, efficient implementation techniques can drastically reduce the computational complexity of this operation, e.g., from 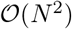 to 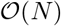, since only the main diagonal elements of **Σ_x_** are required in the update rule, i.e., 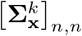. Therefore, the overall computational complexity of the EM algorithm at each iteration is dominated by 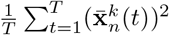, which is a common term in all other MM-based approaches, e.g., convex bounding, MacKay, and LowSNR-BSI. Interested readers can refer to [126] for a computational analysis of EM and other Type-II methods. Figure 8-(B) also depicts the median and inter-quartile ranges of the final negative log-likelihood loss – attained after convergence – of different variants of LowSNR-BSI and Champagne as well as Champagne using EM and MacKay updates.

## Acknowledgment

This result is part of a project that has received funding from the European Research Council (ERC) under the European Unions Horizon 2020 research and innovation programme (Grant agreement No. 758985).

AH acknowledges scholarship support from the Machine Learning/Intelligent Data Analysis research group at Technische Universität Berlin. He further wishes to thank the Berlin International Graduate School in Model and Simulation based Research (BIMoS), the Berlin Mathematical School (BMS), and the Berlin Mathematics Research Center MATH+ for partial support. CC was supported by the National Natural Science Foundation of China under Grant 62007013. GK acknowledges partial support by the Bundesministerium fur Bildung und Forschung (BMBF) through the Berliner Zentrum for Machine Learning (BZML), Project AP4, RTG DAEDALUS (RTG 2433), Projects P1 and P3, RTG BIOQIC (RTG 2260), Projects P4 and P9, and by the Berlin Mathematics Research Center MATH+, Projects EF1-1 and EF1-4. KRM was supported in part by the Institute of Information & Communications Technology Planning & Evaluation (IITP) grant funded by the Korea Government (No. 2017-0-00451, Development of BCI based Brain and Cognitive Computing Technology for Recognizing Users Intentions using Deep Learning) and (No. 2019-0-00079, Artificial Intelligence Graduate School Program, Korea University), and by the German Ministry for Education and Research (BMBF) under Grants 01IS14013A-E, 01GQ1115, 01GQ0850, 01IS18025A, 031L0207D and 01IS18037A; the German Research Foundation (DFG) under Grant Math+, EXC 2046/1, Project ID 390685689. SSN was funded in part by National Institutes of Health grants (R01DC004855, R01EB022717, R01DC176960, R01DC010145, R01NS100440, R01AG062196, and R01DC013979), University of California MRPI MRP-17454755, the US Department of Defense grant (W81XWH-13-1-0494).

